# Brain defence by the extracellular matrix protein Cochlin

**DOI:** 10.1101/2025.11.05.686104

**Authors:** Raghumoy Ghosh, Inyoung Jeong, Preethi Rajamannar, Nathalie Jurisch-Yaksi, Bernett Teck Kwong Lee, Suresh Jesuthasan

**Affiliations:** Lee Kong Chian School of Medicine, Nanyang Technological University, Singapore; Department of Molecular Biology, Umeå University, Sweden; Department of Clinical and Molecular Medicine, Norwegian University of Science and Technology, Trondheim, Norway

## Abstract

The vertebrate brain is protected from infection by tight barriers. However, several barrier structures, including circumventricular organs, can be breached by pathogens. Here, we show that Cochlin, an extracellular matrix-binding protein with anti-bacterial properties, is produced in barrier structures and contributes to immune defence. Transcriptome analysis and *in situ* hybridisation indicate that *cochlin* is expressed in the pineal gland, area postrema, choroid plexus and discrete regions of the meninges of zebrafish, mice and humans. The protein is present in the cerebrospinal fluid, and on the surface of ventricle and meninges. *Cochlin* expression increases in the brain of infected animals and patients, and delivery of recombinant Cochlin reduces bacterial load in zebrafish infected with *Mycobacterium marinum*. Mutation of *cochlin* inhibits clearance of bacteria from the brain of zebrafish, and this is reversed by supplying the LCCL domain of Cochlin. Barrier tissues thus contribute to brain defence by secreting Cochlin.

## Introduction

Infections of the brain present significant clinical challenges due to their complex aetiology, varied presentations, and potential for severe morbidity and mortality. Aside from innate and adaptive immune cells, defensive elements include physical structures such as the meninges, blood-brain barrier and blood-cerebrospinal fluid barrier, which are normally sealed by epithelial junctions. The barriers have a number of potential weak points, however. The circumventricular organs such as the pineal gland and area postrema, which are characterised by their periventricular location, have fenestrated capillaries that facilitate the release of hormones from the brain and sampling of blood-borne substances (1,2). Another barrier structure, the choroid plexus, can become permeable upon infection (3). Here, we identify a new feature of immune defence at brain barriers, that acts independently of anti-microbial neuropeptides (4) and tightening of junctions (5).

## Results and Discussion

To explore immune mechanisms that might counter entry of pathogens via circumventricular organs, we examined a single cell transcriptome of the pineal gland obtained from adult zebrafish (6). We found that a cluster with exosome-producing cells, which are defined by *asip2b* (7), expresses immune related genes such as *cxcl18b, myd88* and *cxcl14* (Fig. S1). This cluster also expresses *cochlin* (Fig. 1a), which was identified originally for its role in progressive deafness but subsequently shown to have diverse functions including stem cell proliferation, otolith crystallization in the ear, mechanosensing in the eye, and innate immunity in the spleen and ear (8–12). Cochlin can bind to the extracellular matrix (ECM), including to collagen VII (13), via two von Willebrand domains (14). The N-terminal *Limulus* factor C, Cochlin and LgL1 (LCCL) domain, which is released by endopeptidase-mediated cleavage, can aggregate bacteria, stimulate the release of pro-inflammatory cytokines, and attract and activate neutrophils and macrophages (9,10,15). *Cochlin* thus represents a candidate component of defence at brain barriers.

**Fig. 1.**
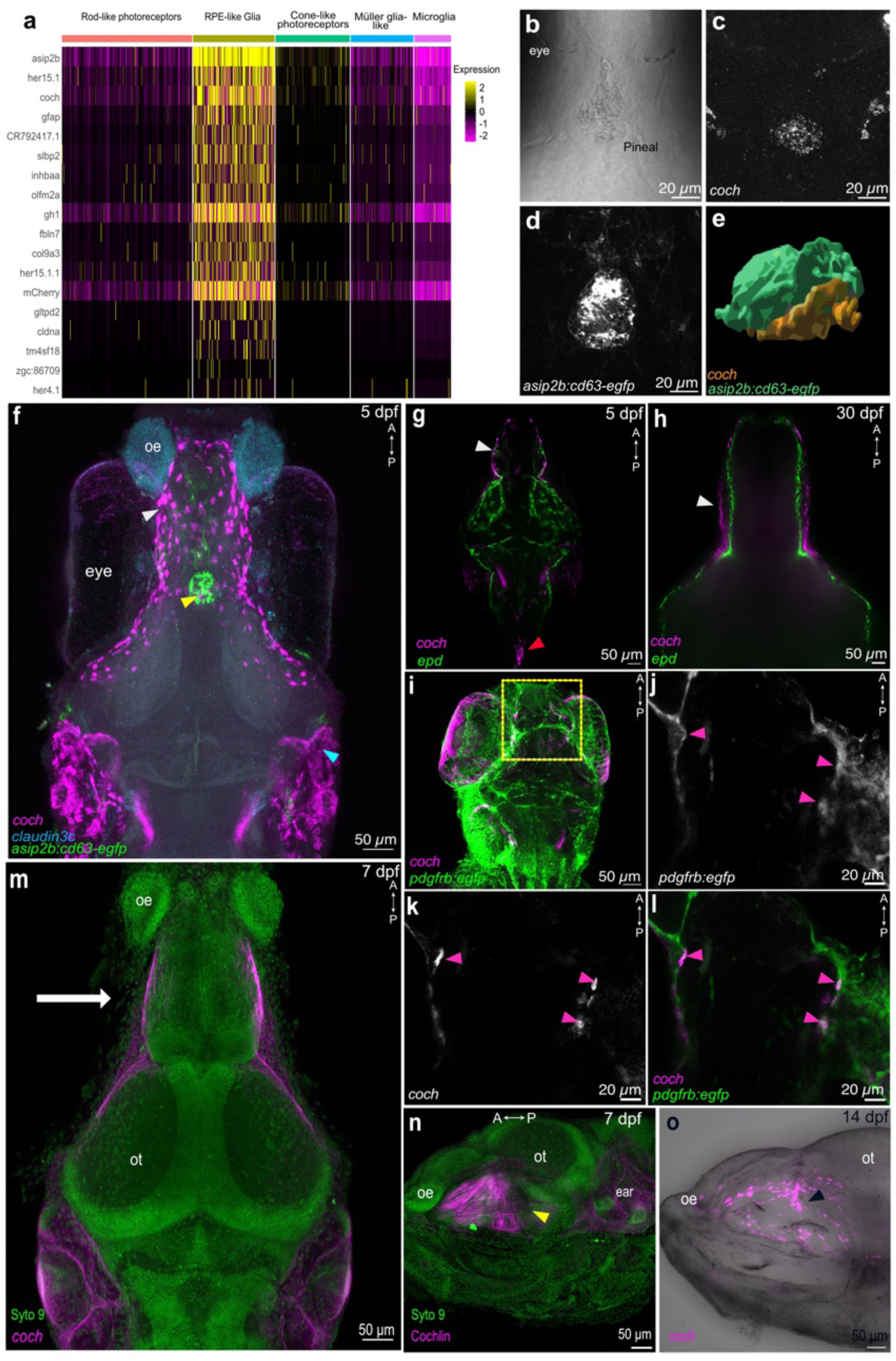
*coch* expression in the pineal gland and meningeal cells of larval zebrafish. **a.** Heatmap of genes expressed in the pineal gland. **b-d**. Dorsal view of the pineal gland of a 7dpf *Tg*(*agrp2:gal4, uas:cd63-egfp*) larvae stained for *cochlin* (HCR-ISH). **b.** Transmitted light image; **c.** *coch* mRNA; **d.** CD63-EGFP protein; **e.** 3D volume rendering of *coch* ISH stain and eGFP fluorescence of *asip2b* positive cells, showing posterior view. **f**. HCR-ISH of *coch* and *cldn3c* in 7dpf Tg(*agrp2:gal4, uas:cd63-egfp*) larvae. Arrowheads showing different localisation of *coch* expression: white – meningeal expression, yellow – pineal expression, and cyan – otic expression. **g.** HCR-ISH of *coch* and *epd* in 5dpf larvae. Arrowheads showing different localisation of *coch* expression: white – meningeal expression, red – area postrema expression. **h.** A single slice from the head of a 30 dpf juvenile zebrafish following HCR-ISH of *coch* and *epd.* White arrowhead shows the layer of the meninges expressing *coch*. **i.** Reperesentative z stack of HCR-ISh for coch in 4 dpf Tg(pdgfrβ*:gal4, uas:egfp*).**j,k,l.** Single slice of the region denoted by the yellow square in (i) showing coch expressing cells (magenta arrow heads). **m, n**. Localisation of Cochlin protein in a 7 dpf larvae, seen in dorsal (m) and lateral (n) views. The white arrow in (m) shows the direction of imaging for (n). Cochlin is present in fibrils in the lateral surface of the brain, medial to the eyes (yellow arrow head, n). **o.** Lateral view, showing *coch*-expressing cells (black arrowhead) medial to the eye in 14 dpf juvenile zebrafish. The eyes have been removed in the samples in panels (m-o). oe: olfactory epithelium; ot: optic tectum.

To determine the spatial distribution of *cochlin* transcripts, hybridization chain reaction - fluorescent in situ hybridisation (HCR-FISH) (16) was performed. Expression was detected in the ventral part of the pineal gland (Fig. 1b-e), i.e. close to the ventricle, in larvae aged 4 days post fertilization (dpf) and older. *Cochlin* transcripts were also detected in the meninges (Fig. 1f) and in the area postrema - another circumventricular organ (Fig. 1g) from at least 5 dpf onwards (see also Fig. S2a, b, d, e & Fig. S7e-h). Meningeal expression was detected first at the lateral edge of the brain at 2 dpf, with a progressive expansion in the expression (Fig. S3). Signal was detected lateral to the fore- and mid-brain, but not dorsally (Fig. 1g, h). Ventral label was detected in the forebrain, from the mouth to the pre-optic area. Additionally, meningeal expression was detected in a discrete band at the boundary of the brain and spinal cord in 14 dpf and 30 dpf fish (Fig. S2b-c,e-f). To confirm that *cochlin* is expressed in the meninges, samples were co-labelled for *ependymin (epd)* and *insulin like growth factor binding protein 2a (igfbp2a)*, both of which are expressed in the leptomeninges (17). *Cochlin* expressing cells were predominantly exterior to the leptomeninges (Fig. 1g,h, Fig. S4). Thus, *cochlin* is expressed in at least two circumventricular organs as well as parts of the meninges.

To identify meningeal cells that express *coch*, we examined single cell transcriptome datasets. In the Daniocell dataset (18), which covers early development, *coch* was detected in the broad “mesenchymal cell” cluster (Fig. S5b), including in ‘meningeal fibroblasts (Fig. S5e) (18). Expression in this population was first detectable at 60-70 hours post fertilization (hpf) while the highest fold change was seen at around 108 hpf onwards (Fig. S5f) (18). This is consistent with the HCR-FISH data (Fig. S3). Approximately 6% of cells within the meningeal fibroblast cluster expressed *coch*. In an adult zebrafish meninges scRNA dataset, *coch* has been identified as a differentially expressed gene (DEG) in fibroblasts, including those that express *pdgfrβ* (17). HCR-ISH for *coch* in 4dpf *TgBAC(pdgfrb:eGFP)* larvae showed colocalization of the eGFP signal and the *coch* expression (Fig. 1i-l). Thus, *coch* is expressed a subset of meningeal fibroblasts.

Immunofluorescence with an antibody raised against the C-terminus of the protein (11) indicates that Cochlin is distributed in a fibrillar pattern on the surface in the lateral fore-and midbrain meninges (Fig. 1m, n). Together, these data suggest that *cochlin* is expressed in fibroblasts in the pachymeninges, and the protein is secreted to the outer surface of the meninges.

Another barrier tissue in the brain is the choroid plexus, which generates cerebrospinal fluid (CSF). In zebrafish, the choroid plexuses are located in the dorsal aspects of the forebrain and hindbrain ventricles (Fig. 2a,b) (19). The polarized monolayers of ciliated epithelium of the choroid plexus express *clusterin* (*clu*) and *igfbp2a* (20). Multiplexed HCR-ISH for *coch* and *clu* showed no colocalization of the signals in young fish (Fig. S6a-h). Sparse label was detected in stromal areas of the choroid plexus in older zebrafish (30 dpf and adults) (Fig. 2c-h). Proteomic analysis of CSF extracted from the telencephalic ventricle of adult zebrafish explants show the presence of Coch (Fig. 2 i), at levels that were comparable to retinol binding protein 4 (Rbp4), a retinol transporter, known to be abundant in zebrafish (21) and human (22) CSF, and two additional CSF abundant proteins, prostaglandin D2 synthase b protein (Ptgdsb.1, orthologous to the mammalian L-PGDS) (23), and the teleost specific Ependymin (Epd) (17). These data establish that Cochlin is secreted into the brain ventricle, in addition to being deposited on the surface of the meninges.

**Fig. 2.**
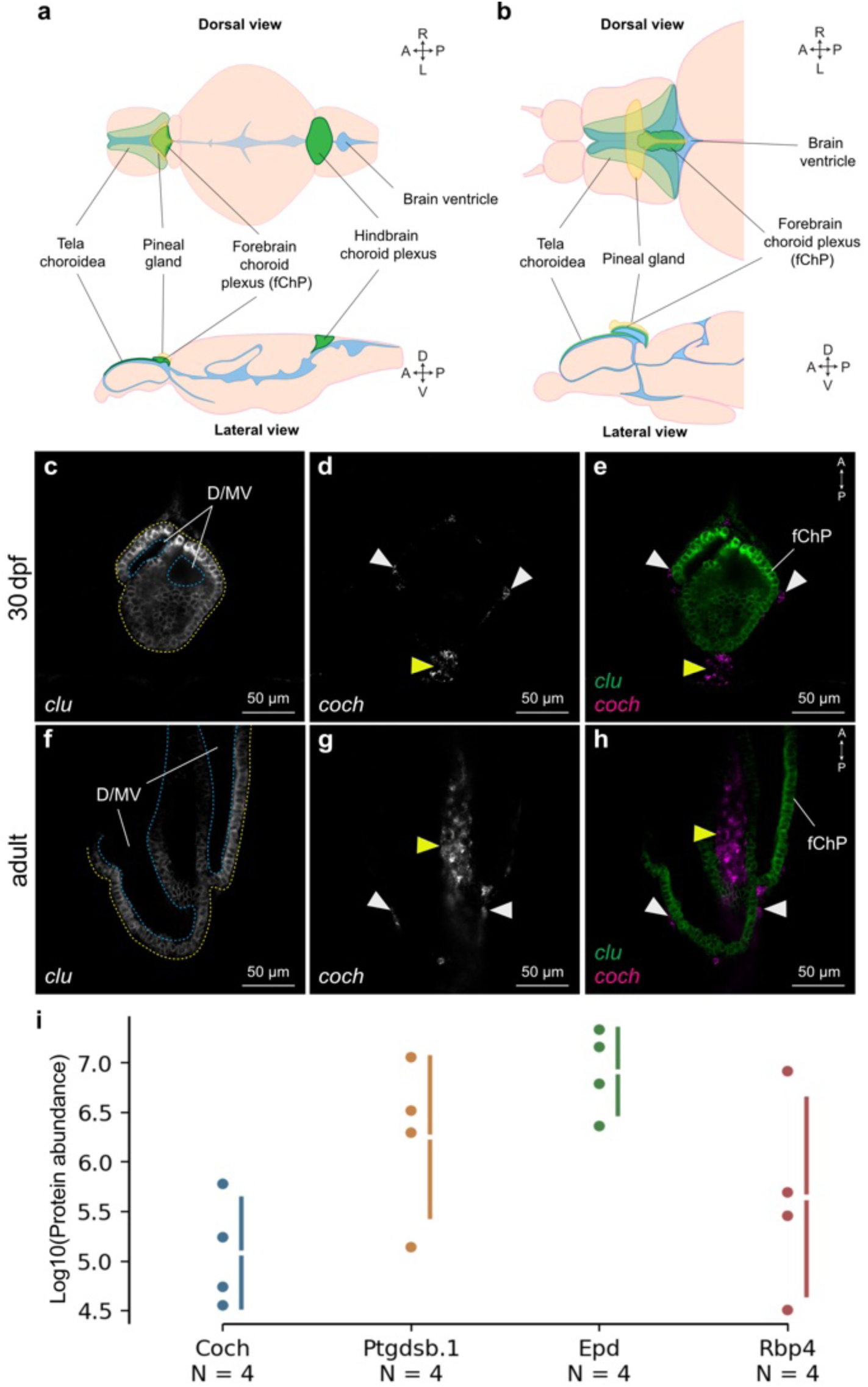
Coch expression in the zebrafish choroid plexus. **a-b**. Schematics of zebrafish brain anatomy in (a) juvenile and (b) adult stages. Green: the choroid plexus, light cream: brain parenchyma, blue: brain ventricles, yellow: the pineal gland. **c-h**. Multiplexed HCR-ISH of *coch* and *clu* mRNAs in (c-e) 30 dpf and (f-h) adult forebrain choroid plexus. Dorsal views of single plane images. Expression of (c, f) *clu* and (d, g) *coch* mRNAs. (e, h) Merged images of each channel. **i.** Coch, Ptgdsb.1, Epd and Rbp4 protein levels in the adult zebrafish telencephalic CSF expressed as log10 values of raw protein reads obtained from LC-MS/MS. Each dot represents one sample of CSF pooled from four fish.

We asked whether expression of *coch* in barrier structures is evolutionarily conserved. Reanalysis of a publicly available integrated day time scRNAseq dataset (24) shows that *Coch* is expressed in vascular and leptomeningeal cells (VLMCs) of the rat pineal gland (Fig. S7). A multi-species database of pineal gene expression indicates that *coch* is produced in the pineal of humans, rhesus monkey, mice and chickens (Fig. S8) (25,26). The Allen Brain Atlas indicates the presence of *Coch* transcripts in the mouse meninges (Fig. S9, S10) (mouse.brain-map.org/experiment/show/71717614) (27). The expression was more prominent in the forebrain meninges, compared to caudal parts of the brain (Fig. S9 a-f; S10a-h).

*Coch* was also detected in the third and lateral ventricle choroid plexus of adult mice (Fig. S10 d-I, S11 d,e) (27–30). This is consistent with a recent transcriptome analysis of these adult tissues, where expression was detected in base barrier cells (31). No expression was detected in the choroid plexus at embryonic stages, based on ISH data in the Allen Brain Atlas and Genepaint (Fig. S11 f-m) (27–30,32). To assess whether *Cochlin* expression in the choroid plexus is developmentally regulated, a snRNA dataset of murine choroid plexus, which included samples from embryonic (E16.5), adult (4 months post-natal) and aged (20 months post-natal) C57BL/6J mice, was analyzed (33). Low resolution clustering was done to identify the main types of cells of the choroid plexus, i.e. epithelium, endothelium, mesenchymal and immune cells, as well as glial and neuronal cells (Fig. S12a). *Coch* was found to be a differentially expressed gene for mesenchymal subset (Fig. S12c). To correlate with the data described above in Fig. S11 with the lack of *Coch* expression in the embryonic choroid plexus, the dataset was split by sample age. *Coch* expression was found in the adult and aged choroid plexus, but not in the embryonic samples (Fig. S13b).

To determine the localisation of Cochlin protein, antibody labelling was performed on adult mice brains, using a previously validated monoclonal antibody (9,10). The brains were split along the midline, to expose the ventricular surface (Fig. 3a), prior to labelling. Cochlin was detected within cells of the choroid plexus of the third and fourth ventricles (Fig. 3b-e). Additionally, Cochlin was detected in a fibre-like pattern in the ventricular wall (Fig. 3f). In the same samples, the lateral leptomeninges near the junction of the olfactory bulb (OB) and the cortex (C) (Fig. 3 a, g, i) showed punctate labelling, as well as intracellular label within fibroblasts. Imaging of the lateral leptomeninges over the caudal cortex (Fig. 3h) showed only puncta. This distribution pattern is consistent with the expression zones detailed in Fig. S9-11.

**Figure 3:**
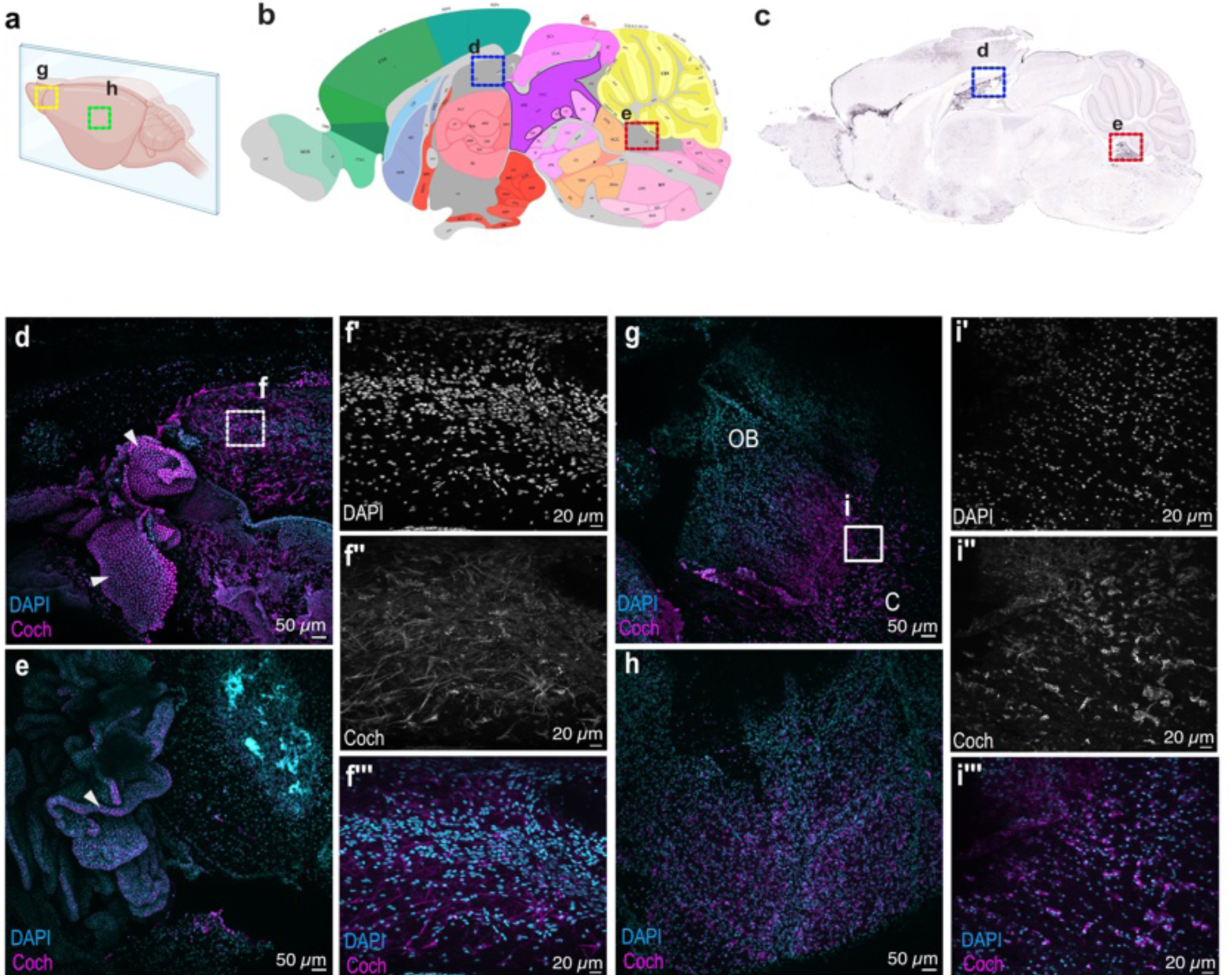
Coch protein localization in the adult murine brain. **a.** A illustration of the brain, showing the plane at which the brain was split prior to antibody labelling. **b.** Sagittal view of the mouse brain at the midline, showing the areas imaged in panels d and e. CTX - cortex, ORB - Orbital area, PL- Prelimbic area, V3- third ventricle, V4- fourth ventricle, TH- thalamus, CBX- cerebellum. Allen Reference Atlas – Mouse Brain (29). **c.** Expression of *Coch* in an adult mouse brain (ISH) at the same sagittal plane as (b). Allen Mouse Brain Atlas, mouse.brain_map.org/experiment/show/71717614 in the forebrain (27–30). **d.** Coch protein localization in the choroid plexus of the third ventricle (arrowheads), as detected by immunofluorescence. The area imaged is indicated by the blue box panels b and c. **e.** Coch protein localization in the choroid plexus of the fourth ventricle (arrowhead); the imaged area is denoted by the red box in panels b and c. **f’-f’’’.** Higher magnification of the third ventricle wall, in the area indicated by the white dashed line box in (d). Cochlin protein is **g.** Immunofluorescence staining for Coch in the lateral leptomeninges at the junction of the olfactory bulb (OB) and cortex (C). Imaging area denoted as the yellow dashed line box in (a). **h.** Immunofluorescence staining for Coch in the lateral leptomeninges on distal cortical region. Imaging area denoted as the green dashed line box in(a). **i’-i’’’.** Higher magnification of Coch immunofluorescence at the junction of the OB and C. Imaging area denoted as the white solid line box in (**g**).

Given the role of Cochlin in aggregating bacteria and attracting innate immune cells, and the increased susceptibility of *Coch^-/-^*mouse to infection in the ear (10) as well as the lung (9), we hypothesized that brain expression of *coch* could influence CNS infections. To assess if infection might induce an increase in *coch* expression in the brain, we analysed a dataset that compared single nuclear transcriptomics of all meningeal cells in an early post-natal *E.coli* meningitis mouse model (GSE221678) (34). *Coch* transcripts could be detected in multiple cell types including pial fibroblasts in both the integrated control and infected datasets (Fig. S14c-d). Visualization of the transcripts in the Violin plot of the merged Seurat object of control and infected datasets shows an increase in the infected dataset clusters (Fig. S15c). Thus, *Coch* may be upregulated in meningitis in mice.

To determine whether infection can affect *cochlin* expression in other barrier tissues in mammals, a human snRNA dataset (35) was examined. GSE159812 contains snRNA sequences from 30 frontal cortex and choroid plexus samples from 14 control individuals (with 1 terminally ill influenza patient) and 8 COVID-19 patients. The highest amount of *COCH* transcripts was detected in the epithelium and mesenchymal cell clusters in the overall dataset (Fig. 4a). Reanalysis and visualization of gene expression levels of the choroid plexus in the COVID-19 patients and the non-viral controls show a higher expression and larger number of cells expressing *COCH* in the COVID-19 group (Fig. 4b-d; unpaired median difference between Control and Covid-19 = 0.434 [95.0%CI 0.185, 0.694]). Mean sample level COCH expression was also higher in the ChP of COVID-19 patients (Fig. 4e-f; unpaired Hedges’ g between Control and Covid groups = 1.08 [95.0%CI 0.154, 2.12]).

**Figure 4:**
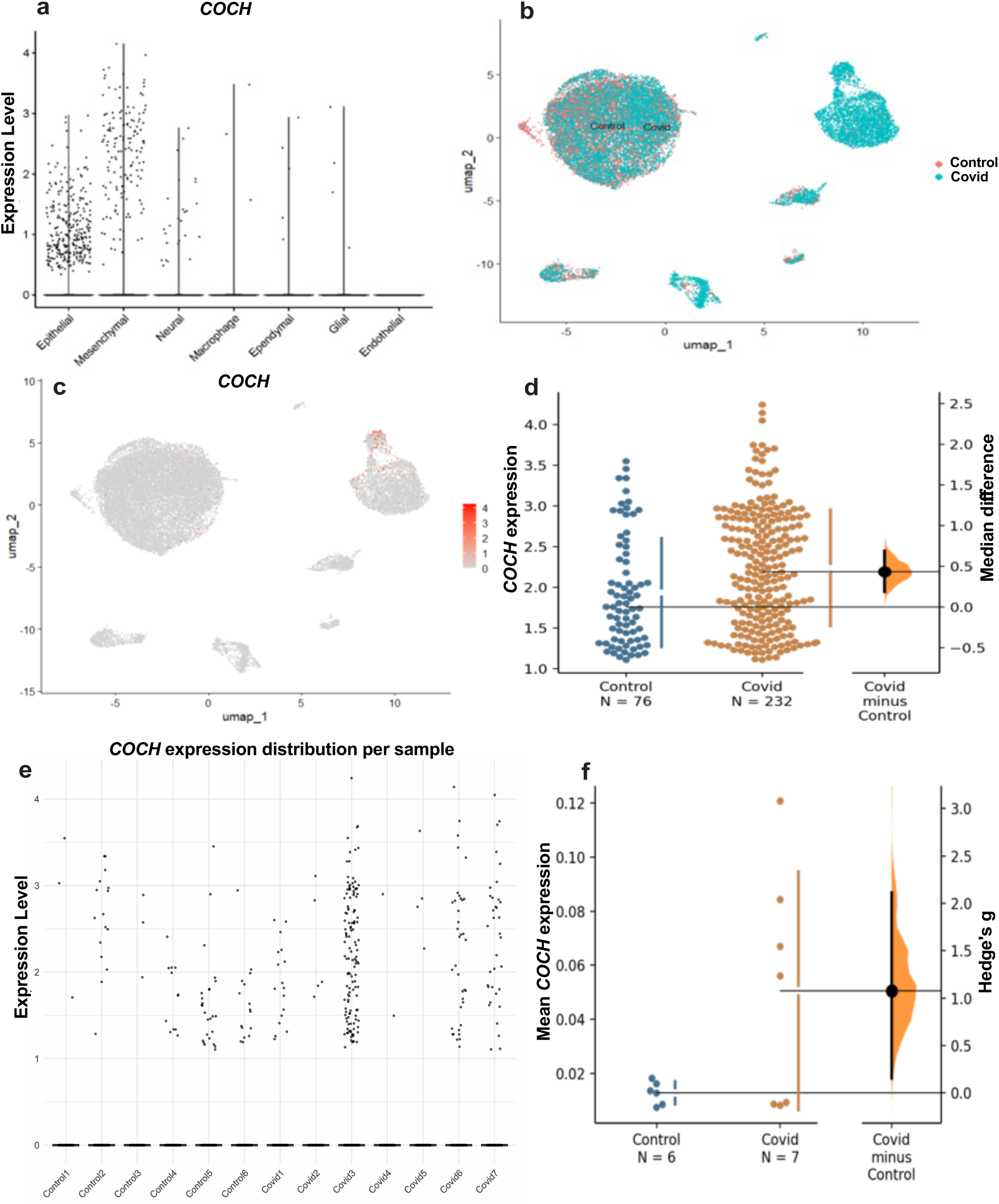
*COCH* expression in the mammalian brain. **a.** Aggregated gene expression plot for *COCH* expression in the cumulative (controls and COVID 19 infected) human choroid plexus dataset (32). The plot is available in the online database (https://twc-stanford.shinyapps.io/scrna_brain_covid19/). **b.** Umap of the integrated Seurat object for ChP from the control and COVID-19 infected donors, split by donor group. **c.** UMAP of the integrated Seurat object for ChP from the control and COVID-19 infected donors, overlaid with *COCH* expression. **d.** Gardner-Altman estimation plot of *COCH* expression in control and covid groups, as extracted from the integrated Seurat object for ChP from the control and COVID-19 infected donors. Each data point represents one cell. Mann Whitney test p-value =0.001. **e.** Dot plot showing *COCH* expression per biological replicate, with each column representing a sample and dots indicating individual cell-level expression. **f.** Gardner-Altman estimation plot of sample level mean *COCH* expression in control and covid groups, as extracted from the integrated Seurat object for ChP from the control and COVID-19 infected donors. Each data point represents one sample. Unpaired t test p value =0.061. **g.** Schematic showing midline sectioning (as denoted the blue plane) of adult murine brain. **h.** Anatomical annotations of expected midlined sectioned surfacea. CTX - cortex, ORB - Orbital area, PL- Prelimbic area, V3- third ventricle, V4- fourth ventricle, TH-thalamus, CBX- cerebellum. Allen Reference Atlas – Mouse Brain (29).

We hypothesized that exogenous Cochlin, delivered to the brain, would inhibit bacterial infection in zebrafish larvae. To test this, we used *Mycobacterium marinum*. Due to its low pathogenicity to humans, smaller replication time and genetic similarity to *M. tuberculosis*, *M. marinum* has been used to model tuberculosis in zebrafish (36). Parenchymal or hindbrain ventricular injections has been used to model brain infection of tuberculosis (37). While Cochlin’s innate immune role has been demonstrated against murine models of both gram positive and gram negative bacteria (9,10), no study has assessed its anti-mycobacterial properties. Furthermore, tubercular meningitis represents a severe healthcare challenge (38).

Recombinant Human COCH showed no direct effect on in vitro culture of *M. marinum* within the range tested (0.08 – 40 μg/ml) (Fig. 5a). Intraventricular injection (1 nl of 40 μg/ml) at 3 dpf did not alter viability of the larvae compared to controls when followed up to 4 dpi (Fig. 5b). 3 dpf wild type larvae (n = 120) were then infected via intraventricular injection with 1 nl of *M. marinum* single cell suspension. Retrospective CFU counting showed an average of 500 CFUs were injected into each larva. 15 larvae were injected with media as injection control. 50% of the infected larvae were randomly selected for intraventricular injection of 1 nl of HuCOCH (40 μg/ml in PBS) 2 hours after bacterial infection and are hereby referred to as the treated group. The remaining infected larvae were injected with the vehicle and this group is referred to as the infection control group. Larvae that died or showed any oedema by the first 24 hours of injection were excluded from the study, as these are unlikely to be specific to the pathogen. 41 infection control larvae and 60 treated larvae were monitored for mortality and humane endpoints for 4 days (39). The treated group had a lower mortality (p value = 0.09, hazard ratio (Mantel-Haenszel) =-0.48) (Fig. 5 c). At 4dpi, fluorescence pixel counting (FPC) was performed, and total area of fluorescent pixels were compared between the infected control and the treated larvae. The treated group had a lower FPC compared to the infection control group (Fig. 5 d-f; unpaired median difference between Control and Treated is -5.99e+03 [95.0%CI - 9e+03, -9.8e+02).

**Figure 5.**
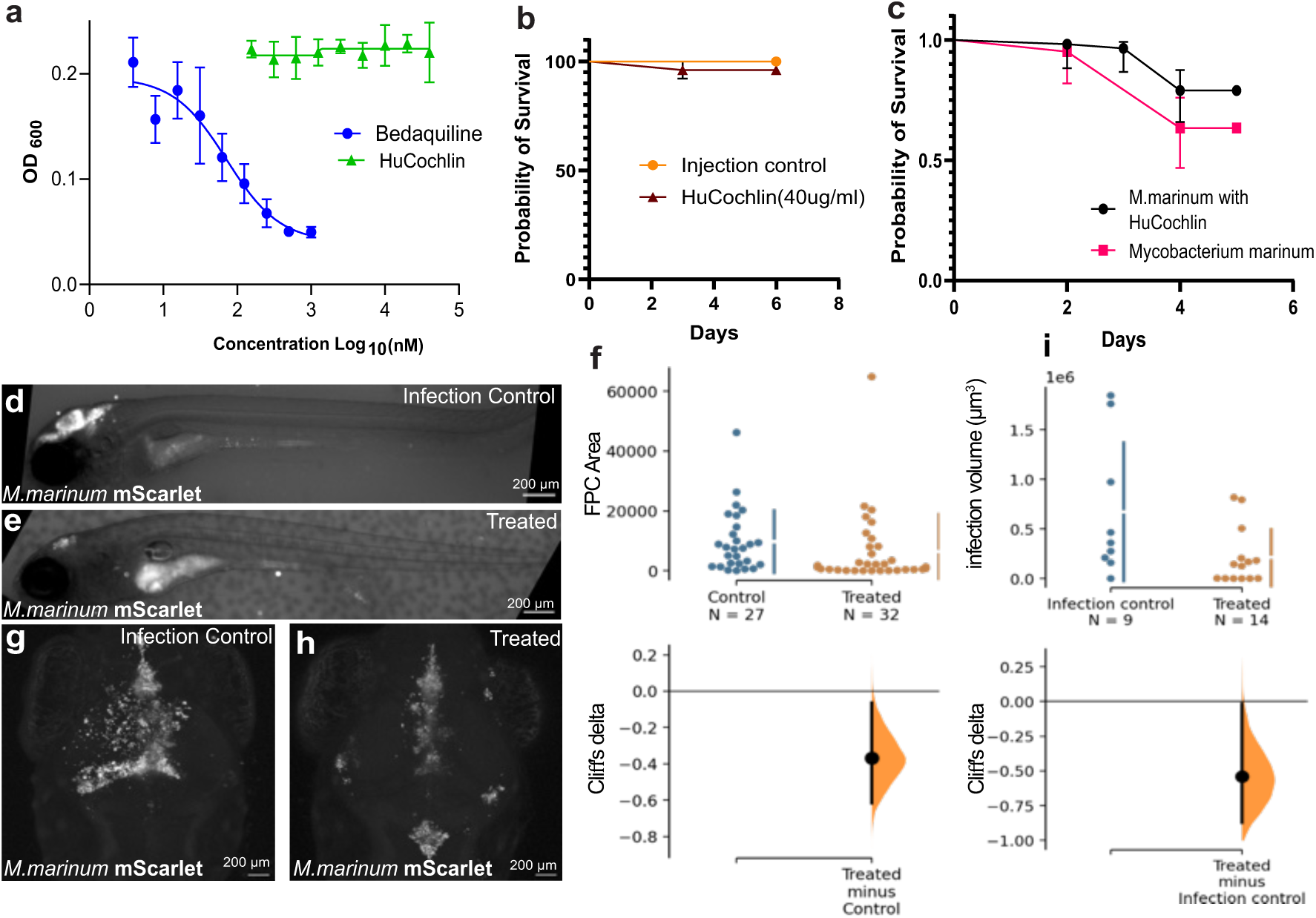
The effects of exogenous Cochlin on bacterial burden in *M. marinum* infected zebrafish larvae. **a.** Minimum inhibitory concentration assay with HuCochlin and Bedaquiline (positive control). **b.** Kaplan Meyer curve for survival estimation between injection control (PBS) and HuCochlin injected 3 pdf larvae. **c.** Kaplan Meyer curve for survival estimation between infection control (500 CFU of *M. marinum*) vs treated group (1 nl of 40 μg/ml HuCochlin in PBS + 500 CFU of *M. marinum*). **d, e.** Representative images of infection control (e) and treated (f) larvae. **f.** The median difference between Control and HuCoch, shown in a Gardner-Altman estimation plot. Both groups are plotted on the left axes; the mean difference is plotted on a floating axis on the right as a bootstrap sampling distribution. The median difference is depicted as a dot; the 95% confidence interval is indicated by the ends of the vertical error bar. Mann Whitney test p-value = 0.03. **g,h.** Representative z-stacks of infection control (g) and treated (h) larvae fixed at 4 dpi and imaged at 20x magnification. **i.** Gardner-Altman estimation plot of infection volume in the infection control and treated groups. Unpaired t test p-value =0.03.

As an alternative method of assessing the effect of Cochlin on CNS bacterial burden, the volume of infection was measured. In a separate experiment, 30 wildtype larvae were injected with 500 CFU of *M. marinum* at 3 dpf, and half of them were treated with intraventricular injection of HuCochlin (1 nL of 40 μg/ml) within 2 hours of the bacterial injection. After exclusion of dead/dying larvae, 9 larvae from the infection control group and 14 from the treated group were fixed at 4 dpi and the cranial region was imaged by confocal microscopy. The treated group had smaller infection volume (unpaired Hedges’ g between Control and Treated is -0.916 [95.0%CI -1.81, -0.0604]) in the imaged region compared to the infection control group (Fig. 5g-i). These data suggest that Cochlin can reduce the spread of *M. marinum* when delivered to the brain.

We next asked whether loss of *coch* increases the spread of *M. marinum* in the brain, using a line generated by Crispr/Cas9 mutagenesis (Fig. 6a). The isolated allele, *coch^lkcsj1-/-^*, had two deletions (Fig. S16b,c): a 132 bp region that included the signal peptide and splice donor of exon 3, and an 8 bp deletion in exon 4. No signal was detected with immunofluorescence in homozygous mutants (Fig. 6c, d), suggesting a loss of the protein. To test the effect of this mutation on response to infection, 144 larvae from in-cross of F1 heterozygous fish were infected at 3 dpf by hindbrain ventricular injection of 1 nL *M. marinum*. Retrospective CFU counting showed that on average 200 bacteria were injected. The larvae were imaged at 4 dpi, then genotyped by PCR. After exclusion of unsuccessfully genotyped samples a total of 135 larvae were used for FPC analysis. *coch^lkcsj-/-^* larvae had higher FPC volume than both the heterozygous (*coch^lkcsj1+/-^*) and wildtype (*coch^lkcsj1^*^+/+^) siblings (respective unpaired Cliff’s delta are -0.289 [95.0%CI -0.504, -0.0572] and -0.415 [95.0%CI -0.622, -0.193], Kruskal-Wallis with Dunn’s multiple comparison test p value of 0.036 and 0.0028 respectively) (Fig. 6e-h). To assess whether the effects of Cochlin are mediated by the LCCL domain produced in the brain, a peptide with this domain was injected into randomly selected half of infected mutants in a separate experiment. This led to a reduction in bacterial load (Mann Whitney test p value = 0.02, unpaired Cliff’s delta between Untreated *coch^lkcsj-/-^* and Treated *coch^lkcsj-/-^*is -0.378 [95.0%CI -0.641, -0.0577]) (Fig. 6i). Together, these data suggest that Cochlin can act in the brain, via its LCCL domain, to inhibit *M. marinum*.

**Fig. 6.**
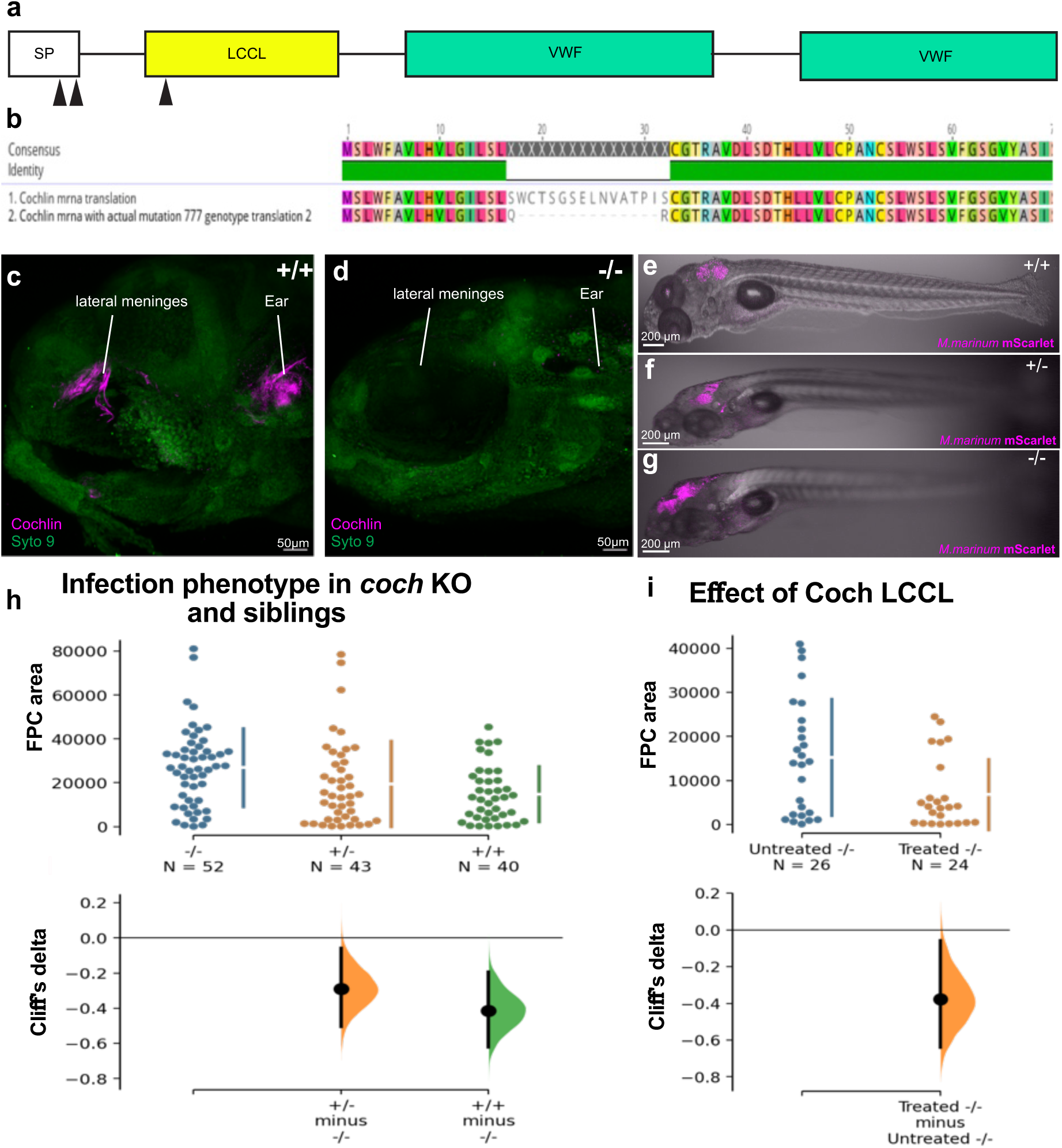
Brain *M.marinum* infection in *coch* mutants. **a.** Schematic showing the target sites (black arrow heads) of the selected gRNAs. **b.** Schematic showing the deletion in the *coch* protein as per the in silico analysis. **c & d.** Representative z-stack images of 5dpf *coch^lkcsj1+/+^*(c) and *coch^lkcsj1-/-^* (d), stained with anti-Coch antibody. **e - g.** Representative images for FPC analysis of *coch^lkcsj1+/+^* (e), *coch^lkcsj1+/-^* (f) and *coch^lkcsj1-/-^* larvae (g) at 4dpi. **h.** Cumming estimation plot with Cliff’s delta for 2 comparisons against the shared control -/-. **i.** Gardner-Altman estimation plot with Cliff’s delta for FPC area between untreated and treated *coch^lkcsj1-/-^* at 4dpi. (+/+ = *coch^lkcsj1+/+^*, +/- = *coch^lkcsj1+/-^*, -/- = *coch^lkcsj1-/-^*).

In summary, these findings establish that Cochlin is produced by structures that form the interface between the brain and the external world, including the pineal gland, area postrema, choroid plexus and meninges. Within the brain, we show that Cochlin is present in the CSF of zebrafish; COCH has recently also been detected in human CSF (40). In the mouse, we show that Cochlin is present in a fibrillar pattern on walls of the ventricle. These data suggest that Cochlin is secreted into the ventricle, where it binds to the extracellular matrix. This enables Cochlin to act on bacteria outside the blood-brain barrier, complementing its previously reported function in the spleen (9). Cochlin is also present outside the brain, at discrete regions of the meninges that are close to openings in the skull. We speculate that the LCCL domain is released in these locations upon infection, causing the aggregation of bacteria while attracting immune cells and triggering the release of pro-inflammatory cytokines IL-6 and IL-1ß (10,15). These data thus suggest a previously unappreciated aspect of brain defence: the secretion of dormant matrix-associated proteins, which can be activated to counter pathogens that have reached the walls of the brain. Previous work has shown that Cochlin can act on *Staphylococcus aureus* and *Pseudomonas aeruginosa* (9,10), and we show that it also affects *Mycobacteria marinum*. It would be informative, in the future, to perform infection experiments on juvenile or adult fish, as expression in the choroid plexus is seen at these stages and this would be predicted to provide increased defence. It would also be important to assess the effects of Cochlin on other pathogens that can infect the brain, such as *Porphyromonas gingivalis* and *Pseudomonas aeruginosa*. If this is effective, an enhancement of mechanism identified here may be a therapeutically useful strategy against a range of pathogens. Finally, an additional implication of the findings here is that in patients with autoimmune sensorineural hearing loss triggered by T-cells reactive to Cochlin (41,42), it would be predicted that there may also be infiltration of pro-inflammatory auto-immune T-cells to the brain. The extent of this remains unknown.

## Acknowledgments

This work was supported by funds from the Ministry of Education, Singapore (RG34/20), the National Research Foundation, Singapore (NRF2017-NRF-ISF002-2676); the Research Council of Norway (RCN FRIPRO grants 353348 to N.J.Y. and NORBRAIN infrastructure grant 295721), and an NTU Research Scholarship to R.G. The spinning disk confocal imaging was performed at the Cellular and Molecular Imaging Core Facility (CMIC), Norwegian University of Science and Technology (NTNU). CMIC is funded by the Faculty of Medicine and Health Sciences at NTNU, Central Norway Regional Health Authority and NALMIN II (NFR 322607).

We thank Simon Larsson of the Phil Ingham Lab, and Samsher Singh of the Kevin Pethe lab, LKC School of Medicine, NTU, for the stocks of *M. marinum* and technical help with zebrafish infections. We thank Fredericus van Eeden for providing the zebrafish Cochlin antibody. We thank Lars Hagen and Animesh Sharma for the proteomics analysis. Proteomics and Modomics Experimental Core Facility (PROMEC) is part of NAPI (RCN INFRASTRUKTUR-program 295910). PROMEC computations are performed on resources provided by Sigma2—the National Infrastructure for High Performance Computing and Data Storage in Norway (Projects NN9036K and NS9036K). We thank Björn Schröder, Héloïse Chat, Sanja Vanhatalo and Sandra Holmberg for providing mouse tissue. We also thank Anna Barron for discussions. Katja Kerner and Sofie Beutels contributed to generating a part of the multiplexed HCR-FISH images under the supervision of Inyoung Jeong and Nathalie Jurisch-Yaksi. The *TgBAC(pdgfrb:eGFP)* line was provided courtesy of Prof. Naoki Mochizuki of the National Cerebral and Cardiovascular Center, Osaka, JAPAN.

## Methods

### Zebrafish lines

Zebrafish were grown and maintained in recirculating fish systems (Tecniplast) on a 14-hour light/10-hour dark cycle. The following fish lines were used: *TgBAC(agrp:Gal4-VP16)^tlv04^; Tg(UAS:EGFP-HsaCD63)^lkc3^*, *TgBAC(pdgfrβ:EGFP)^ncv22T^* (43), *nacre* (44), AB, and *coch^lkcsj1^*.

Animals up to 30 dpf were analysed irrespective of their sex, as this cannot be determined at these stages. Sex distribution for adults used for HCR-ISH are listed in Table S1.

### Hybridization Chain Reaction-Fluorescent In situ Hybridization (HCR-FISH)

The qHCR probes for *coch, foxj1b, clu, igfbp2a, epd, rgra* and *claudinh* were supplied by Molecular Instruments. The ISH was carried out according to the manufacturer’s v10.0 protocol for HCR-RNA- FISH whole mount zebrafish embryos and larvae with modifications as necessary for age of embryo/larvae. For elucidating the expression timeline, wildtype AB larvae were used for 2, 3 and 7dpf, nacre for 5, 14, 30 dpf and adult, *Tg(kdrl:egfp, lyve1b:dsred)* larvae for 5 dpf, *TgBAC(pdgfrb:eGFP)* larvae for 4 dpf and *Tg(asip2b:gal4, uas:cd63-egfp)* larvae for 6 dpf. Proteinase K treatment was titrated based on the age of the larvae, as described previously (45). For the 14, 30 dpf and adult forebrain choroid plexus imaging, the brains were dissected before HCR-FISH, as described previously (19,46). For the 14 dpf (Fig S2 and Fig1h), the entire head was cleared by using the Binaree Tissue ClearingTM Kit (#HRTC-012, Binaree, Republic of Korea) before HCR-FISH, as described previously (47). For the adult samples, both male (n=5) and female (n=7) fish were used and there was no sex difference. Both genders were represented in this study. Labelled samples were imaged using an upright Zeiss Axio Imager LSM800, Examiner LSM880 or Nikon spinning disk (Nikon Ti2-E with CrestOptics X-Light V3) confocal microscopes, using a 20x (0.8 NA) apochromat objective, a 20x Plan-Apochromat (NA 0.8) objective or a Nikon 20x Plan APO (NA 0.8) objective. IMARIS and Huygens were used for the 3D reconstructed images (Fig 3c-d) and deconvolved cross sectional images (Figure 3e-f and FigS2), respectively.

### Antibody labelling- zebrafish larvae

Antibody staining for Cochlin was done as described previously (11,48). In brief, 7 dpf *nacre^-/-^* larvae were fixed overnight in 4% paraformaldehyde (PFA) at 4°C, washed thrice in PBS with 0.1% Tween20 (0.1% PBST) for 10 mins each, and dehydrated in 100% methanol for at least 48 hours. Then the larvae were rehydrated with 0.1% PBST washes. For antigen retrieval, the larvae were incubated in 150 mm HCl of pH 9 for 5 minutes at RT and 15 minutes at 70° C. Then the larvae were washed twice in 0.1% PBST and twice in sterile distilled water before incubating in 100% acetone at -20°C for 20 minutes. Subsequently the acetone was washed out with 2 washes in sterile distilled water and 2 washes in 0.1% PBST, and then the larvae were incubated in the blocking buffer (10% goat serum, 0.8% Triton X, 1% BSA in 0.1% PBST) for 3 hours at 4° C. Post blocking, the larvae were incubated in the antibody solution (1% goat serum, 0.8% Triton X and 1% BSA in 0.1% PBST) in which the anti-Cochlin antibody, described in Leventae et al. and kindly donated by Fredericus van Eeden, was diluted at 1:2000. The larvae were incubated with the primary antibody for 3 days at 4° C on a shaker at 90 rpm. The excess primary antibody was washed out with the following wash buffers: 3×1 hr - 10% goat serum, 1% TritonX in PBS (1% PBSTx); 2×10 minutes - 1% PBSTx, 2×1 hr in 10% goat serum in 1% PBSTx. The larvae were then incubated in the appropriate secondary antibody (AlexaFluor546 goat anti-rabbit) in 1:1000 final dilution, for 4 days at 4° C on a shaker at 90 rpm. The larvae were then washed 3×1 hr in 10% goat serum in 1% PBSTx and 2×1 hr in 0.1% PBST. Before imaging, they were incubated in 300 μL of 0.1% PBST containing Syto 9 for nucleic acid staining (1:5000 dilution of stock concentration; ThermoFisher S34854). The larvae were imaged within a week of staining. Controls which had not been incubated with the primary antibody were used to assess nonspecific binding.

### Antibody labelling- adult murine brain

C57Bl/6J mice were obtained from Charles River Laboratories (Germany), bred inhouse, and housed at 22 ± 1 °C and 55 ± 5 % humidity under a 12-h light/dark cycle in individually ventilated cages in a specific-pathogen-free environment. The freshly dissected whole brains were collected in ice cold PBS from 8 weeks old (n=2) and 13 weeks old (n=3) adult male mice. They were cut along the midline (Fig. 3g), to expose the 3^rd^ and 4^th^ ventricular surfaces and ChPs, and washed in PBS thrice for 10 mins with gentle shaking. Fixation was done with pre chilled acetone at -20 °^C^ for 1 hour followed by 3 x 1hour washes with PBS at 4 °^C^. The samples were then blocked for 3 hours in PBS with 5% BSA. Anti mouse Cochlin antibody (clone 9A10D2, Sigma MABF267) was used at a dilution of 1:1000 in PBS with 5%BSA, for the overnight primary antibody treatment at 4 °^C^ with gentle shaking. The samples were then washed with PBS (1hour, thrice) at 4 °^C^ with gentle shaking. Then they were then incubated in PBS+ 5% BSA with the secondary antibody (goat anti-rat IgG (H+L) cross-adsorbed secondary antibody, Alexa Fluor™ 568 - A-11077,Invitrogen) in 1:1000 final dilution, overnight at 4° C with gentle shaking. After washing with PBS (1hour, thrice) at 4 °^C^ the samples were treated with DAPI for nuclear stain (5mg/ml- 1:5000 dilution in PBS) and imaged.

The samples were imaged with an upright confocal microscope Nikon AX using 16x and 40x water dipping objectives with maximum field of view. The images were analysed in ImageJ (49).

### Bacterial strains and growth conditions

The *Mycobacterium marinum* strain used for this study are of the Aronson strain modified to express the fluorescent protein mScarlet. It was grown at 31°C in Middlebrook 7H9 broth (Difco) supplemented with 0.5% bovine serum albumin, 0.2% dextrose, 0.2% glycerol and 0.3% catalase.

Liquid cultures were prepared with Difco™ Middlebrook 7H9 (Becton Dickinson and Company, USA) medium with 10% Albumin-dextrose-saline (ADS) enrichment with or without 0.5% glycerol (Sigma Life Science). Glycerol was included for culturing conditions and omitted in drug susceptibility assays. Bacteria stocks were banked in 7H9 medium with 15% glycerol and stored in -80°C. Passaging of cultures were limited to twice. For all studies, *M. marinum* was grown to mid-log phase with an OD600 of 0.2-0.8 in Middlebrook 7H9 media (with 10% ADS). Kanamycin was used for selection pressure.

### Determination of minimum growth inhibition concentration

Minimum growth inhibitory concentration was determined via broth microdilution method. Minimum Inhibitory Concentration (MIC50) represents the lowest concentration of compound that results in 50% bacterial growth inhibition. The compounds were tested in triplicates with a concentration range from 0.08 – 40 μg/ml for HuCOCH (Sino Biological, 11368-H07H) and 0.004 – 1 μM for Bedaquilin (BDQ). Inoculums treated with BDQ were used as a positive control. Drug free inoculums with 0.9% DMSO were used as negative control. The HuCOCH protein was diluted to 0.4 mg/ml in Middlebrook 7H9 Broth medium as per the manufacturer’s instructions and added directly to the 96-well clear flat bottom plate. The concentration series of HuCOCH was created by serial dilution. The final OD600 for *M. marinum* was adjusted in Middlebrook 7H9 Broth medium supplemented with 0.05% Tween 80 and 10% ADS enrichment. Thus, 180 of the *M. marinum* culture (OD600 0.0056) with 20 μL of HuCOCH (on respective concentrations) in 7H9 broth were added in each well to maintain final OD600 of 0.005. 200 ml (OD600 0.005) of bacterial culture was then inoculated into each well of the spotted 96-well plates for BDQ. 2 ml (100 times desired concentration) of BDQ in 90% DMSO from the drug intermediate plate was spotted in the respective wells of the 96-well clear flat bottom plates used for determining MIC.

Plates containing *M. marinum* were incubated at 32°C for 72 hours. OD600 was measured using Cytation-3 microplate reader (BioTeck instruments) and dose-response curves were plotted and analysed using GraphPad PRISM 10 (GraphPad Software, USA) from which MIC50 values were obtained.

### Intraventricular injections of *Mycobacterium marinum*

The bacterial processing and injections were done as carried out by Takaki et al., with necessary modifications (50). In brief, *M. marinum* was cultured to mid-log phase with an OD600 of 0.2-0.8 in Middlebrook 7H9 media (with 10% ADS) in a volume of 30 ml. The culture was spun down (3800 rpm for 10 mins) and resuspended in appropriate freezing media (7H9 + 10% OADC +15% glycerol) so that the OD was 4. 200 µl aliquots of 4 OD culture in freezing media were stored at -80°C till the day of infection. On the day of infection, one 200 µL aliquot was first washed twice with 1 ml of sterile 7H9, then spun down and resuspended in 1 ml media. It was then divided into 5 aliquots of 200 µl each. Single cell suspension was created by aspirating and ejecting each aliquot 10 times via a 27-gauge needle. 1 ml of 7H9 + ADC was added to each 200 μL aliquot; these were mixed by inversion or brief vortexing and then centrifuged it at 100 g for 1 min. 1 ml of the supernatant was transferred from each aliquot and the process repeated thrice in total to get 15 ml of supernatant. The collected supernatant was then passed through a 5 µm filter and spun down at 20,000 g for 10 mins. The resultant pellet was resuspended to an OD of 1.6 (standardized to have 300 CFU of *M. marinum* in 1 nl) and then used for injection.

Zebrafish larvae at 3 dpf were anesthetized in buffered tricaine (270 mg/l) and then mounted in 2% low-melting agarose with dorsal side up. A pneumatic microinjector with a fine capillary glass pipette was used to inject 1 nl of processed mycobacterial suspension into the hindbrain ventricle. The larvae were then carefully released and individually housed in 100 μL of E3 media in a 96 well flat bottomed optically clear plate.

### Retrospective Colony Forming Unit (CFU) counting

Retrospective CFU counting of the injection mixture was done to assess number of bacteria injected and consistency of bacterial load delivered into the hindbrain ventricle. 1 nl of the injection mixture was injected into a 200 μL PCR tube with 100 μL of sterile 7H9 broth, via the same needle being used to inject the larvae. Samples were collected after every 1/3^rd^ of total larvae to be injected, thus collecting three samples for every experiment. The samples were then serially diluted to 10 times and 100 times dilutions and plated into 7h11 agar with kanamycin in quadrant plates. The plates were incubated at 30°C and CFU was counted after 10 days. Average CFU counts were reported.

### Mutation of *coch* by CRISPR-Cas9 gene editing

3 targets were first selected in the first 3 exons of Zebrafish cochlin for CRISPR-Cas9 mediated mutagenesis from the CHOPCHOP website (51–53). The resultant crRNA design was checked in the IDT CRISPR design checker tool for availability of pre-designed crRNA. One of the selected targets (cochlin23) was available as a prevalidated crRNA (Dr.Cas9.COCH.1.AD) while the other two (cochlin48, cochlin95) had been amongst the crRNA used by Leventae *et al.* (11) for creating *coch* crispants. The on target (higher value indicates more efficient cutting) and off target scores (higher value indicates less off target effects) as per IDT crRNA design checker has been furnished below. Maximum scores are 100. The gRNA and RNP solutions were created as per the protocol published by Kroll *et al*. (51). F0 bialelic knock outs were created by injecting the RNP solution into the one cell stage of AB eggs and grown up. The resultant F0 larvae were then allowed to grow till 3 months of age and genotyped using the genotyping primers described below. The candidate F0 adults were outcrossed with wildtype AB adults. The resultant F1 heterozygous mutants were selected by genotyping.

crRNA sequences with the on and off target scores.

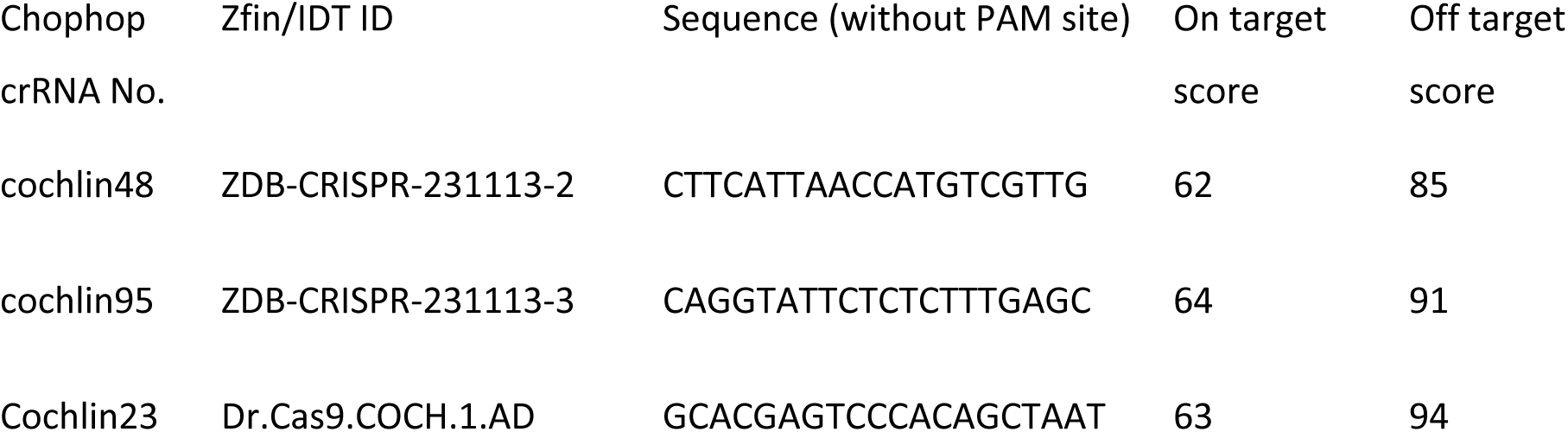

Genotyping of adults and larvae

Caudal fin clipping was used to collect biopsies from individual adult zebrafish for genotyping. For larval genotyping, the whole larvae were taken for genotyping after overdosing with tricaine. In case of IHC validation of the stable *coch* KO in F2 generation larvae obtained from the in-cross of F1 heterozygous (*coch*^+/-^) parents, mini-fin clips from individual larvae were collected. The rest of the larva was collected in individual PCR tubes for PFA fixation and IHC processing. The DNA was extracted using alkaline hotshot method (54). The primers used for identification of *coch* KO mutants are: Forward primer TGGTGCCTGGTAAAATGCTCT; Reverse primer TACCTGTGAATGGCTGCTCC. Standard 20 μL PCR reactions with Taq polymerase and annealing temperature of 62°C was used to amplify the 922 bp (wildtype) region and gel electrophoresis was done to identify mutants.

### Fluorescent Pixel Counting and analysis

The assessment of bacterial burden in larvae at 4 dpi was done by fluorescent pixel counting (FPC) as published previously (50), with minor modifications. In brief, the larvae were washed thrice in sterile E3 media with tricaine (270 mg/l). They were then individually transferred to a 96 well flat bottomed optically clear plate with 100 μL of E3 media and incubated on ice for 30 mins to ensure that they sink to the bottom. Imaging was carried out in the LSM800 inverted confocal microscope with 2.5x objective. 3-5 uninfected control larvae were included in the imaging for threshold calculations. After imaging, the larvae were either used for post infection CFU enumeration or genotyping. The images were analyzed in ImageJ using the method and FPC macro described by Takaki et al (50).

### Imaging of heads of infected larvae

Infected and control larvae were first anesthetized with an overdose of tricaine and then fixed overnight in 4% PFA. The fixed larvae were washed in PBS and then mounted in 1.5 % low melting temperature agarose in E3 medium. All Z-stacks were collected on Zeiss LSM800 upright confocal microscope with a 20X objective. Images were analyzed in ImageJ (49) using the 3D object counter plugin (55) and the cumulative volume of fluorescent objects were taken to assess the bacterial burden.

### Analysis of pineal gland single cell RNA sequencing data

The datafiles corresponding to the sample “GSM3511192” were used in this analysis (6). The data was processed using the Seurat R package v5.1.0 (56). Quality control parameters of unique feature count between 200-2500 and mitochondrial transcripts less than 5% were used to filter out doublets, low quality and dying cells. Post normalization and scaling per the Seurat commands, principal component analysis (PCA) was done by Seurat’s “RunPCA” command. Number of dimensions for the dataset was assessed to be 10 by elbow plot. Clustering was performed by graph-based (K nearest neighbor) clustering algorithm (Seurat’s “FindNeighbours” and “FindClusters”). Dimension reduction was done with Seurat’s “RunUMAP” with same number of dimensions used for PCA analysis. Markers for the clusters were derived by the “FindAllMarkers” command and the result was compared to existing literature to assign cluster identities. Cluster 1 was reassigned as RPE-like glial cells on the basis of differential expression of *agrp2*. Specific markers for RPE-like glial cells were then derived by “FindMarkers” with log average fold change filter of 0.25 and ROC test. Genes with an average log2 fold change greater than 2 were selected for downstream analysis.

### Gene Ontology Annotation and Filtering

Selected marker genes were annotated using the clusterProfiler package (v4.6.0) and the org.Dr.eg.db annotation database. GO terms were retrieved via the bitr() function, and biological process (BP) terms were isolated for functional enrichment. To identify immune-related processes, a recursive search was performed starting from the parent GO term GO:0002376 (“immune system process”) using the GOBPCHILDREN mapping from the GO.db package. All descendant immune-related GO terms were compiled. To refine immune gene annotations, the Ensembl BioMart interface (biomaRt v2.54.0) was queried using the drerio_gene_ensembl dataset. Genes associated with the compiled immune-related GO terms were retrieved using getBM() with attributes including Ensembl gene ID, external gene name, GO ID, and ontology namespace.Genes from the pineal dataset annotated to immune-related GO terms were filtered and deduplicated. The final list of immune-associated genes was visualized using the DoHeatmap() function in Seurat, with feature labels suppressed for clarity (label = FALSE).

### Visualization of Daniocell dataset

The Daniocell dataset (18) was visualized by using the RDS file “Daniocell2023_SeuratV4.rds” and “cluster_annotions.csv” available at https://daniocell.nichd.nih.gov/index.html. The overall dataset was subsetted for mesenchymal cells based on the tissue annotations. The mesenchymal Seurat object was further split by the “meningeal fibroblast” value in the “identity.sub” column of the metadata. Dimplot, FeaturePlot and DotPlot functions were used for data visualization.

### Analysis of published single cell RNA sequencing data rat pineal

The raw data files for rat pineal was available at GSE115723 (24). The data was processed using the Seurat R package v5.1.0 (56). The daytime datasets with extended references (GSM3188344-day1_extendedRef, GSM3188346_Day2_ExtendedRef) were loaded and combined using the “merge” function. Quality control parameters of unique feature count more than 400, ncount_RNA < 20000 and mitochondrial transcripts less than 25% were used to filter out doublets, low quality and dying cells. The combined dataset was then scaled and normalized. Principal component analysis (PCA) was done by Seurat’s “RunPCA” command. Clustering was performed by graph-based (K nearest neighbor) clustering algorithm (Seurat’s “FindNeighbors” and “FindClusters”). Dimension reduction was done with Seurat’s “RunUMAP” with same number of dimensions used for PCA analysis. Markers for the clusters were derived by the “FindAllMarkers” command and the result was compared to existing literature to assign cluster identities. The Dotplot function was used to map relevant genes to the combined dataset.

### Analysis of published transcriptomic datasets of murine meninges

The published single nuclear transcriptomics of all meningeal cells in an early post-natal *E.coli* meningitis model (GSE221678) was similarly processed (34). The two replicates of control mice and three of the infected mice were respectively merged, subsetted with the following QC parameters: unique features > 200 and < 2500, transcript < 6000 and > 500 and mitochondrial gene percentage < 1; and integrated, using “HarmonyIntegrate” method, to integrate the layers into two respective Seurat objects. The control and infected integrated objects were then individually scaled and normalised. After clustering, cluster annotations were assigned as per the DEGs of each cluster and the markers published by the same authors (34). Then the two objects were merged and infected cell cluster were compared to their control counterparts via the “FindallMarkers ()” function.

### Analysis of published single nuclear RNA sequencing data of murine choroid plexus

The published snRNA sequencing dataset on murine choroid plexus across embryonic, adult and aged stages, available at NCBI GEO ID-GSE168704, was re-analysed with Seurat R package without the splitting the dataset by the ages (33). QC parameters used were unique features between 200 and 5000 and mitochondrial gene percentage < 2. The Seurat object was then scaled and normalised with respect to the percentage of mitochondrial genes, clustered with resolution of 0.1. Dimensional reduction was carried out using the “umap” method and 5 dimensions. Clusters were annotated into broad clusters as per the markers published by the same authors (33). DEGs were identified by using the “FindallMarkers ()” function. Visualization of Coch levels was split by the age sub stages by using the VlnPlot function and splitting the object by the “orig.ident” column of the metadata.

### Analysis of published single nuclear RNA sequencing data of human ChP

The human choroid plexus snRNA dataset is available at NCBI GEO ID-GSE159812 (35). The ChP specific datasets for the controls (n=6 : GSM4848456, GSM4848457, GSM4848458, GSM4848459, GSM4848460 and GSM4848461) and the COVID-19 infected donors (n=7, GSM4848451, GSM4848452, GSM4848453, GSM4848454, GSM5171535, GSM5171536 and GSM5171537) was re-analysed with Seurat R package (v5). The single datset concerning ChP in a patient with influenza was left out of the analysis. Raw count matrices were read using the Read10X() function from the Seurat package (v5), and Seurat objects were created with a minimum threshold of 200 detected genes and 3 cells per gene. Mitochondrial gene content was quantified using PercentageFeatureSet() with the pattern ^MT-, and cells with >5% mitochondrial reads, 2500 detected features, or >5000 total counts were excluded to remove low-quality or stressed cells. Filtered samples were annotated with metadata including sample identity (sampleId) and condition (group: Control or Covid), and then merged into a single Seurat object. Post data normalization, scaling and dimensionality reduction, harmony integration was applied to account for inter-sample variability. Differential gene expression between COVID-19 and control samples was assessed using the Wilcoxon rank-sum test via FindMarkers() (log fold change threshold = 0.25, minimum expression in 10% of cells). The identity class was set to group prior to testing. COCH expression was specifically examined using FetchData() and visualized with violin plots, UMAP overlays, and sample-level boxplots. To validate cell-level findings and avoid pseudo-replication, COCH expression was aggregated per sample using group by() and summarise() from the dplyr package, and mean sample level COCH expression values were extracted for comparison.

### Extraction and proteomics of adult zebrafish cerebrospinal fluid

Adult *nacre* zebrafish were euthanised in an ice bath and brains were dissected out in ice cold aCSF (36.208g NaCl, 19.820g D+Glucose, 0.746 KCl, 1.972g MgSO_4_.7H_2_O, 0.851g KH_2_PO_4_, 10.080g NaHCO_3_, 1.470g CaCl_2_.2H_2_O in 5l H_2_O, osmolarity 300-310mOsm) with the meningeal layer intact (46). CSF was manually drained from the telencephalic ventricle by inserting a fine glass capillary into the anterior telencephalic commissure between the two olfactory bulb lobes. An Eppendorf CellTram oil microinjector apparatus connected to the capillary was used to suction the CSF and the liquid was transferred to a cryo-vial containing 5 µL sterile PBS. CSF from four adult fish were pooled together for each replicate and flash frozen in liquid nitrogen. Four replicates containing CSF from four adult fish were resuspended in 100 µL 1% sodium deoxycholate, 100 mM Tris-hydrochloride (pH 8.5), 10 mM Tris(2-carboxyethyl) phosphine, and 40 mM Chloroacetamide. The samples were sonicated and incubated with 0.5 µg trypsin for digestion overnight at 37°C. Peptides were desalted, dried, and resuspended in 0.1% formic acid before performing liquid chromatography with tandem mass spectrometry (LC-MS/MS) using the Bruker Daltonics timsTOF Pro with the nanoElute liquid chromatography system. Peptides were separated for 75 minutes using a 150 µm*25 cm Pepsep 25 (Bruker Daltonics) with running buffers (A) 0.1% formic acid and (B) 0.1% formic acid in acetonitrile with a gradient of 2 - 40% operated in the DIA PASEF mode. Source conditions were as follows: capillary voltage, 1,600 V; dry gas, 3.0 L/min; dry temperature, 180 °C. MS1 and MS2 spectra were acquired within an m/z range of 100–1,700, with collision energy linearly interpolated between 59 eV (1/K₀ = 1.60 V·s/cm²) and 20 eV (1/K₀ = 0.6 V·s/cm²).

Raw MS/MS data was processed using DIA-NN (57) performed against the Uniprot zebrafish proteome library. Oxidation of methionine and acetylation of protein N-terminal as dynamic post-translational modification was appended with maximum allowed variable modifications to 3 per peptide spectrum match. Maximum allowed miss-cleavage were set to 2 and both the MS and MS/MS mass-accuracy was set to 20 ppm. QuantUMS (high accuracy) quantification strategy with protein inference and match-between-runs (MBR) functionality was used to reduce missing values in raw data. Intensity values for each protein group were normalized using the DIA-NN label-free quantification algorithm. Quality control check was done with an R-script (https://raw.githubusercontent.com/animesh/scripts/ccc63e663e0cd905a4325327292ae8f15c1d5885/geneGroupsQC.r).

### Statistics and reproducibility

Statistical analyses were performed in GraphPad Prism 10.3. Shapiro-Wilks test was performed to check for normality. Unpaired t test was performed for data deemed to follow Gaussian distribution as per the results of the Shapiro-Wilks test. Mann Whitney test was performed for comparisons amongst two groups with non-Gaussian distribution. Effect size was calculated as Hedge’s g or median difference or Cliff’s delta using https://www.estimationstats.com/ (58). Sample size calculation for infection experiments was based on difference of means derived from FPC values of infection in F0 knockout for *coch* which have not been shown in this manuscript.

## Code and Data availability

All associated codes will be available upon publication. Images and data used for plotting charts will be deposited on Figshare (10.6084/m9.figshare.32447016).

## Animal ethics

All experiments in Singapore were performed in accordance with protocols A20038 and A21062, which were approved by the IACUC of Nanyang Technological University. The animal facility and maintenance of the zebrafish in Norway were approved by the Norwegian Food Safety Authority. The murine samples were sourced from the Björn Schröder Lab, UMU (Dnr A8-2025). All the procedures here were performed in accordance with the European Communities Council Directive and the Norwegian Food Safety Authorities.

