## Supplementary material for "Brain defence by the extracellular matrix protein Cochlin": (Fig. S1)

### Supplementary Figures

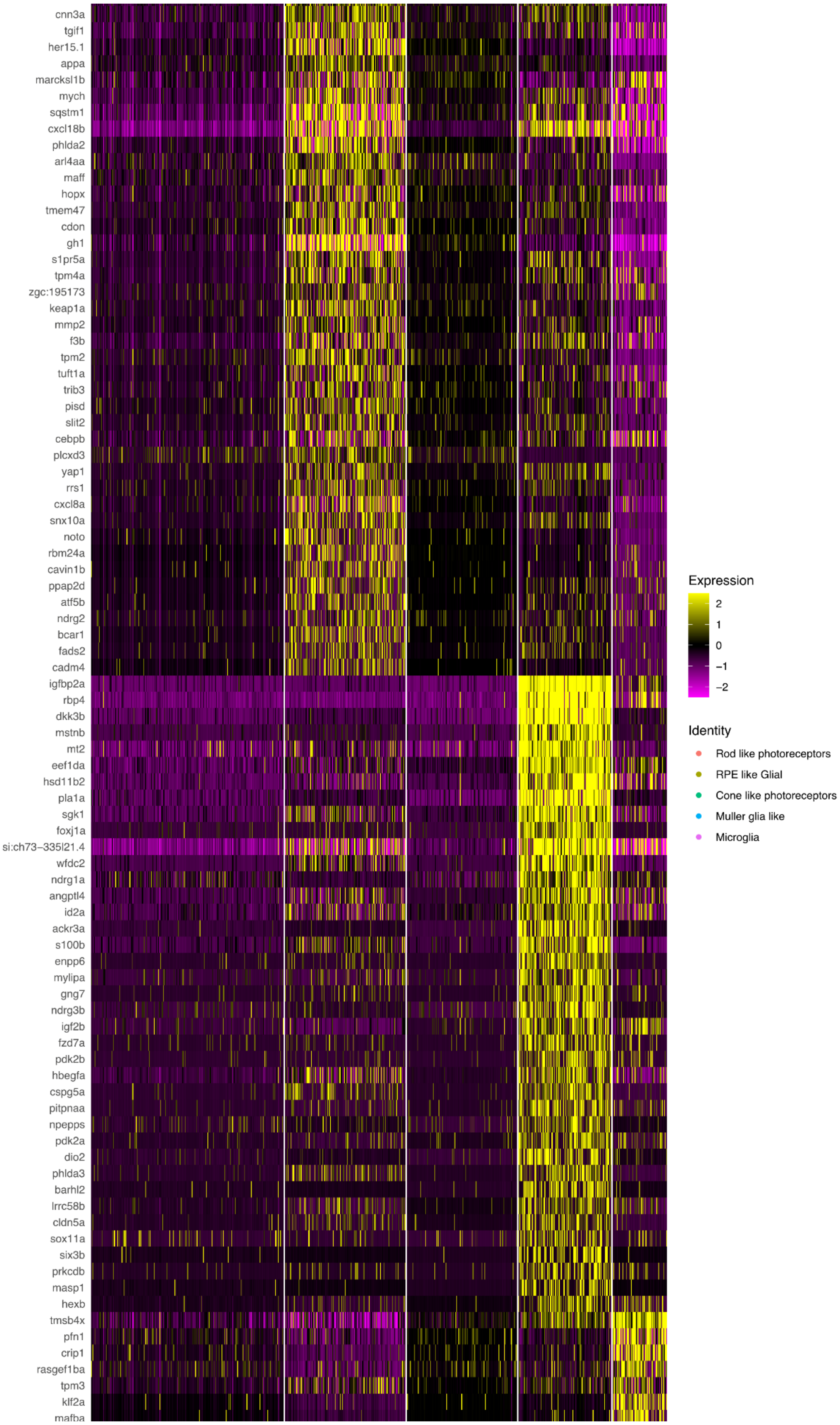

**Figure S1. Expression of immune genes in the zebrafish pineal gland.** Heatmap of immune related genes filtered from the cluster markers of the adult zebrafish pineal gland (1).

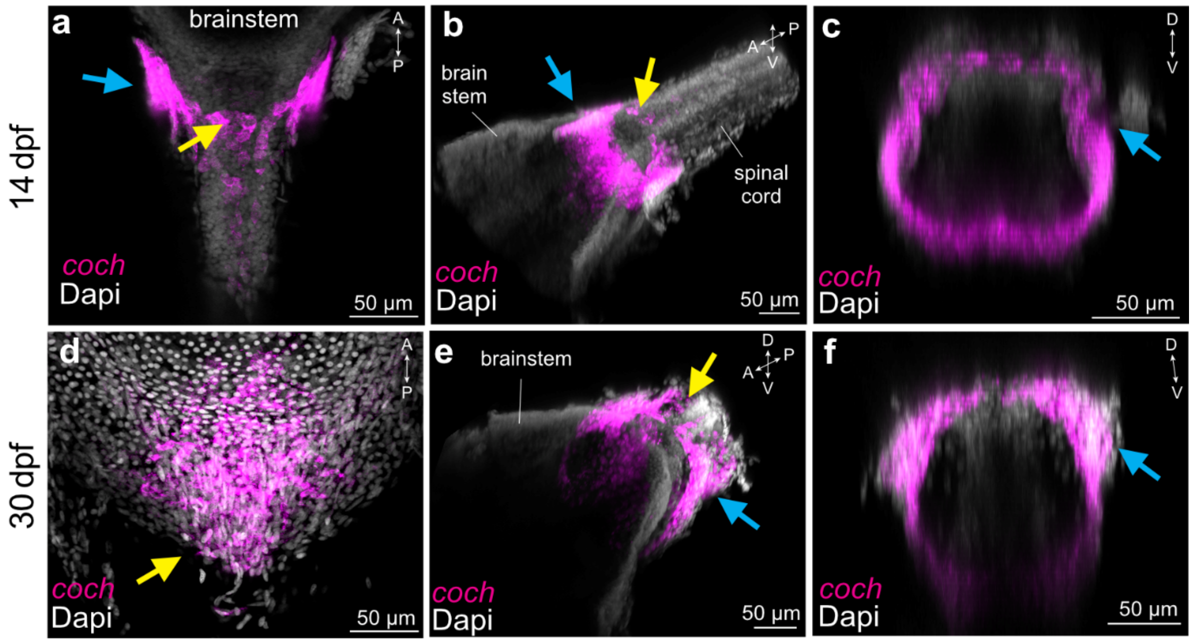


**Figure S2. *coch* expression in the area postrema and around the posterior brainstem of zebrafish of 14 and 30 dpf** . **a-f**. HCR-ISH of *coch* mRNA in the brainstem region at (a-c) 14 dpf and (d-f) 30 dpf. *Coch*: magenta, DAPI: grayscale. Yellow dotted lines: basal sides of the choroid plexus epithelial cells that stromal cells are located. Yellow arrows: meningeal expression of the posterior part of the area postrema. Blue arrows: meningeal expression of a discrete band at the boundary of the brain and spinal cord. **a,d.** Dorsal views of max projection images. **b,e**. 3D reconstructed images of the brainstem region. **c,f**. Cross sectional view of the band-like expression in (b, e). A: anterior, P: posterior, D: dorsal, V: ventral.D/MV: Diencephalic/mesencephalic ventricle.

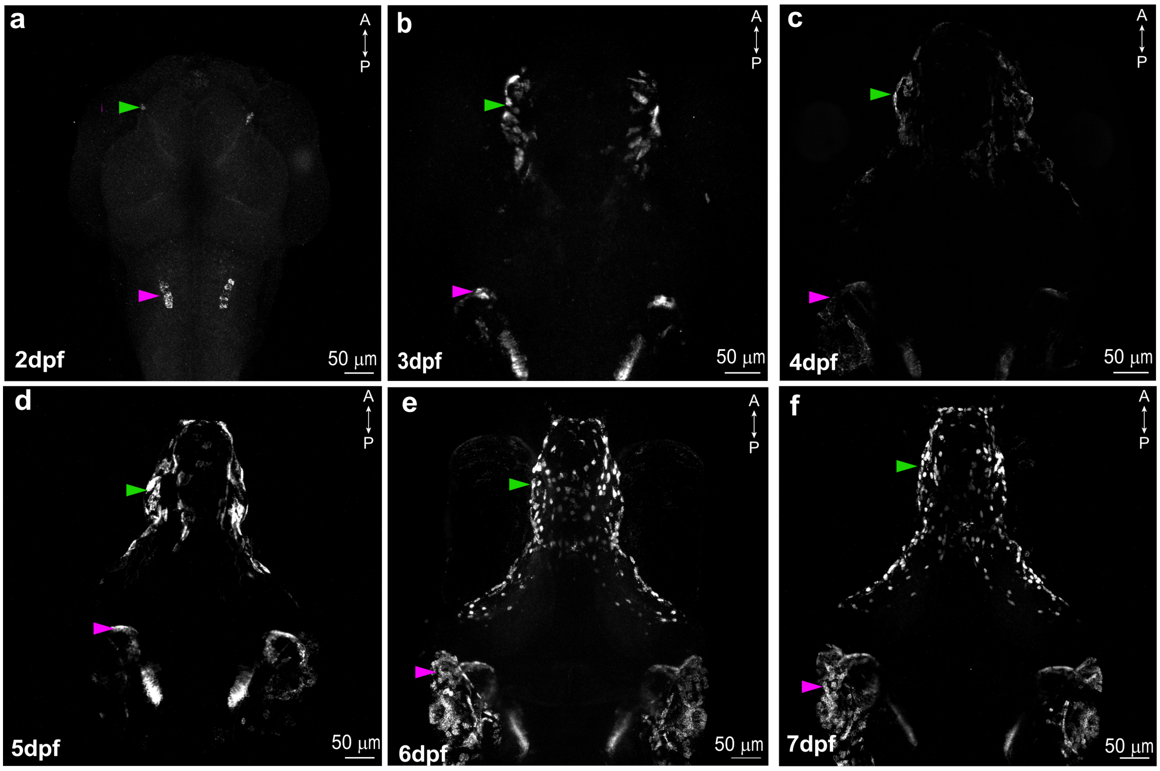


**Figure S3. Time series of *coch* expression in the fore- and midbrain of zebrafish larvae up to 7 dpf**. **a.** 2 dpf, **b.** 3 dpf, **c.** 4 dpf **d.** 5 dpf, **e.** 6 dpf and **f.** 7 dpf. Arrow heads: green – meningeal expression, magenta – otic expression. Expressing cells are located laterally, and then ventrally to the brain.

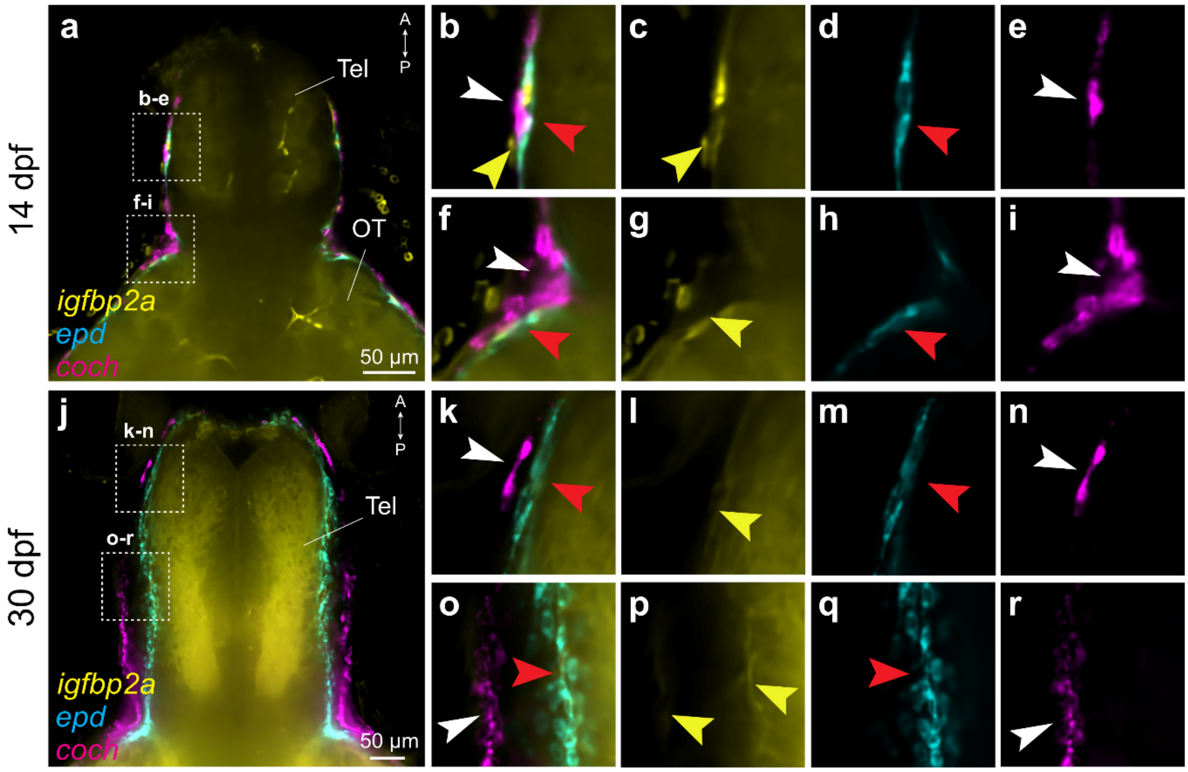

**Figure S4. Meningeal *coch* expression with meningeal markers in the meningeal areas of the zebrafish forebrain in 14 and 30 day old zebrafish larvae**. **a,j.** Multiplexed HCR-ISH of *coch, epd* and *igfbp2a* mRNAs in (a) 14 dpf and (j) 30 dpf. Dorsal views of single plane images. *coch*: magenta, *igfbp2a*: yellow, *epd*: cyan. Merged channels. White dotted boxes show the location of insets b-e, f-i, k-n and o-r. Tel: Telencephalon, OT: optic tectum. **b-e, f-i**. High magnification images of the white dotted boxes in (a). Merged (b,f) and individual (c&g, d&h and e&i) channels. **k-n, o-r**. High magnification images of the white dotted boxes in (j). Merged (k,o) and individual (l&p, m&q and n&r) channels. White arrowheads: the *coch* positive meningeal cell layer, yellow arrowheads: the *igfbp2a* positive meningeal layer, red arrowheads: the *epd* positive meningeal layer. A: anterior, P: posterior, Tel: telencephalon, OT: optic tectum.

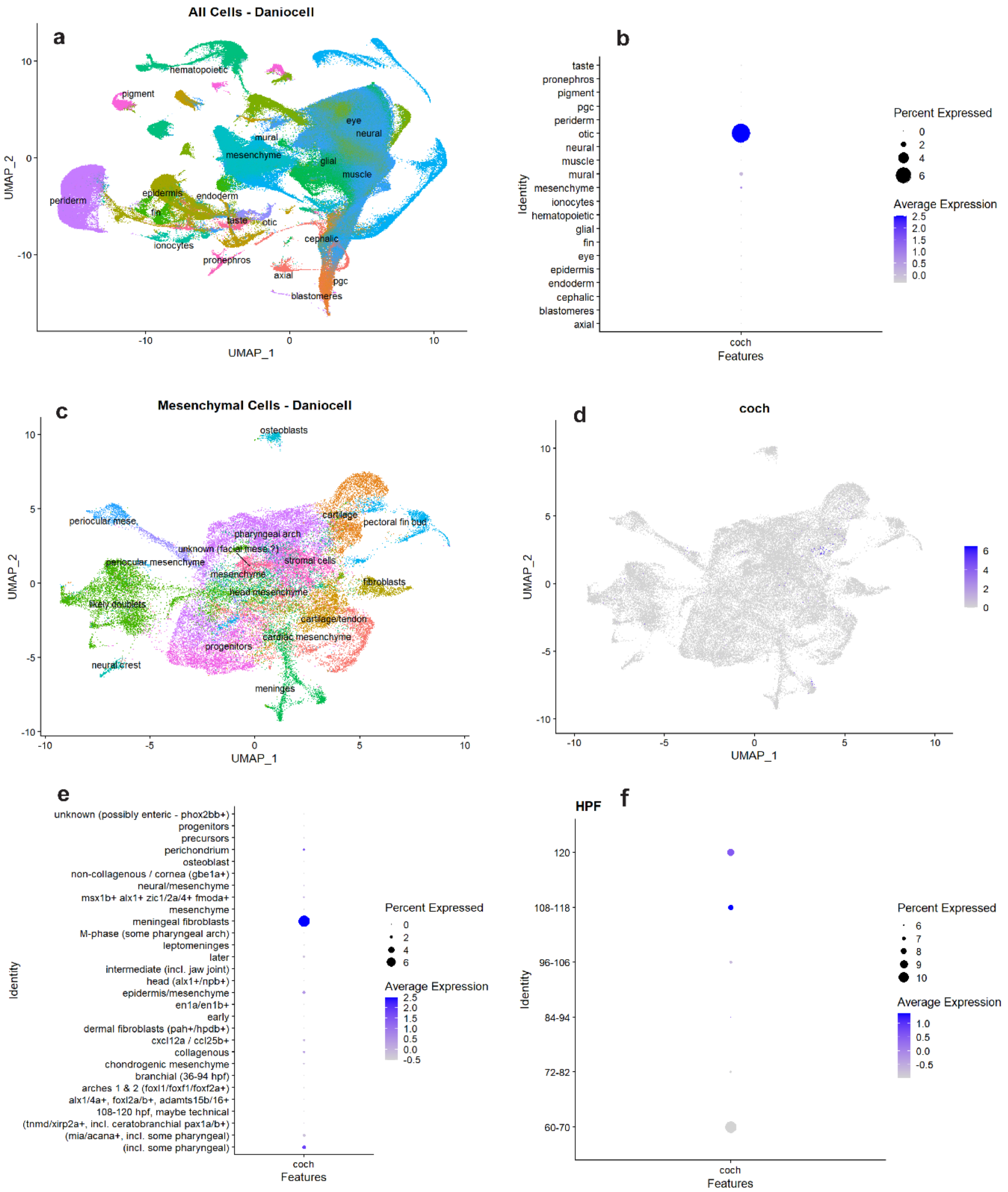


**Figure S5. Identification of cells expressing *coch* in the meninges of larval zebrafish. a.** UMAP of the Daniocell dataset containing all cells. **b.** Dot plot for *coch* expression within the Daniocell dataset showing highest expression in the otic tissue followed by mural and mesenchymal tissue subsets. **c.** UMAP of mesenchymal tissue cell which have subsetted out of Daniocell dataset. **d.** Feature plot for *coch* expression within the mesenchymal tissue subset showing expression in cartilage and meninges subgroup of mesenchymal cell. **e.** Dotplot for *coch* expression within the mesenchymal tissue subset showing highest expression within the meningeal fibroblast cluster. **f.** Dotplot for *coch* expression within the meningeal fibroblasts split by hours post fertilization (hpf) stages.

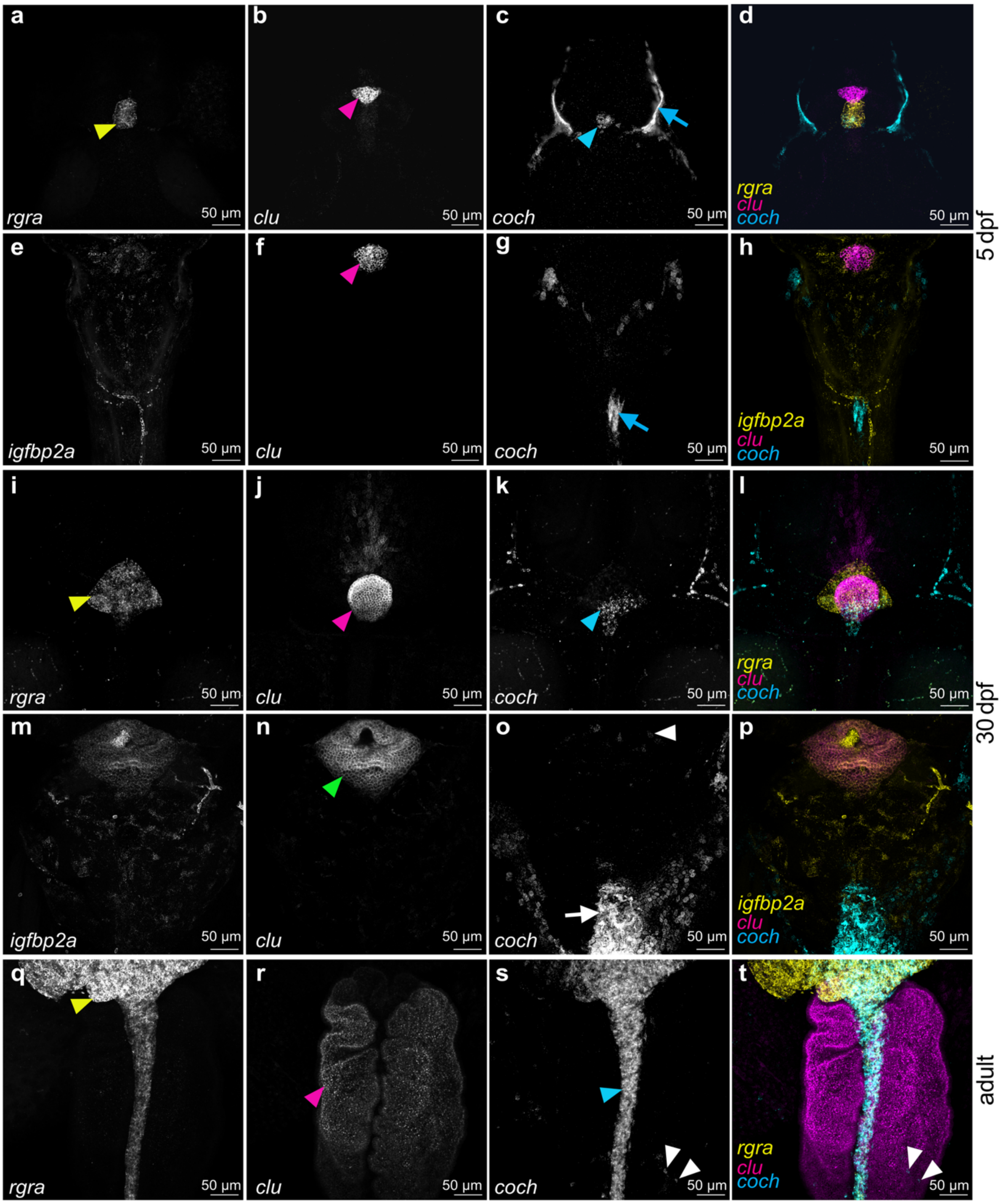


**Fig. S6. Coch expression in the choroid plexus. a-t**. Multiplexed HCR-ISH of *coch*, *igfbp2a*, *clu* and *retinal G protein coupled receptor a (rgra)* mRNAs in **(a-h)** 5 dpf, **(i-p)** 30 dpf and **(q-t)**

adult fish. All images are maximum projections of z-stacks, showing dorsal views. Expression of **(a, i, q)** *rgra* (yellow arrowheads: the pineal gland), **(e, m)** *igfbp2a* **(b, f, j, n, r)**, *clu* (magenta arrowheads: forebrain choroid plexus epithelial cells, green arrowheads: hindbrain choroid plexus epithelial cells), **(c, g, k, o, s)** *coch* (blue arrowheads: pineal gland, white arrowheads: choroid plexus stroma, blue arrows: forebrain meninges, white arrows: area postrema. **d, h, l, p, t.** merged images of each channel. A: Anterior, P: Posterior. Gamma of panels g, k, o and s has been set to 0.45 for better visualization.

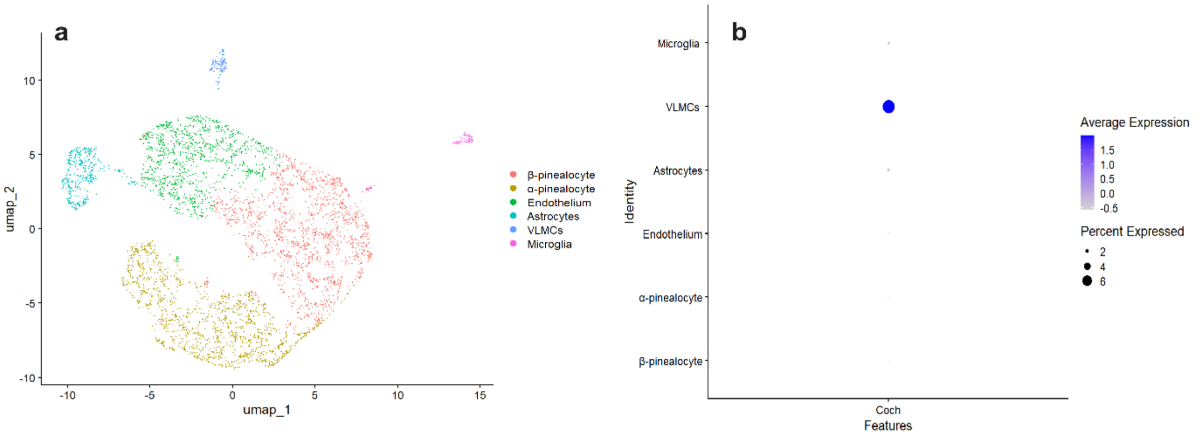


**Figure S7. *Cochlin* expression in the rat pineal gland. a.** UMAP of integrated day time datasets of rat pineal scRNAseq (3). **b.** Dotplot of *Coch* expression as obtained from the reanalysis of published data (3) and integrating day time specific datasets of rat pineal scRNAseq.

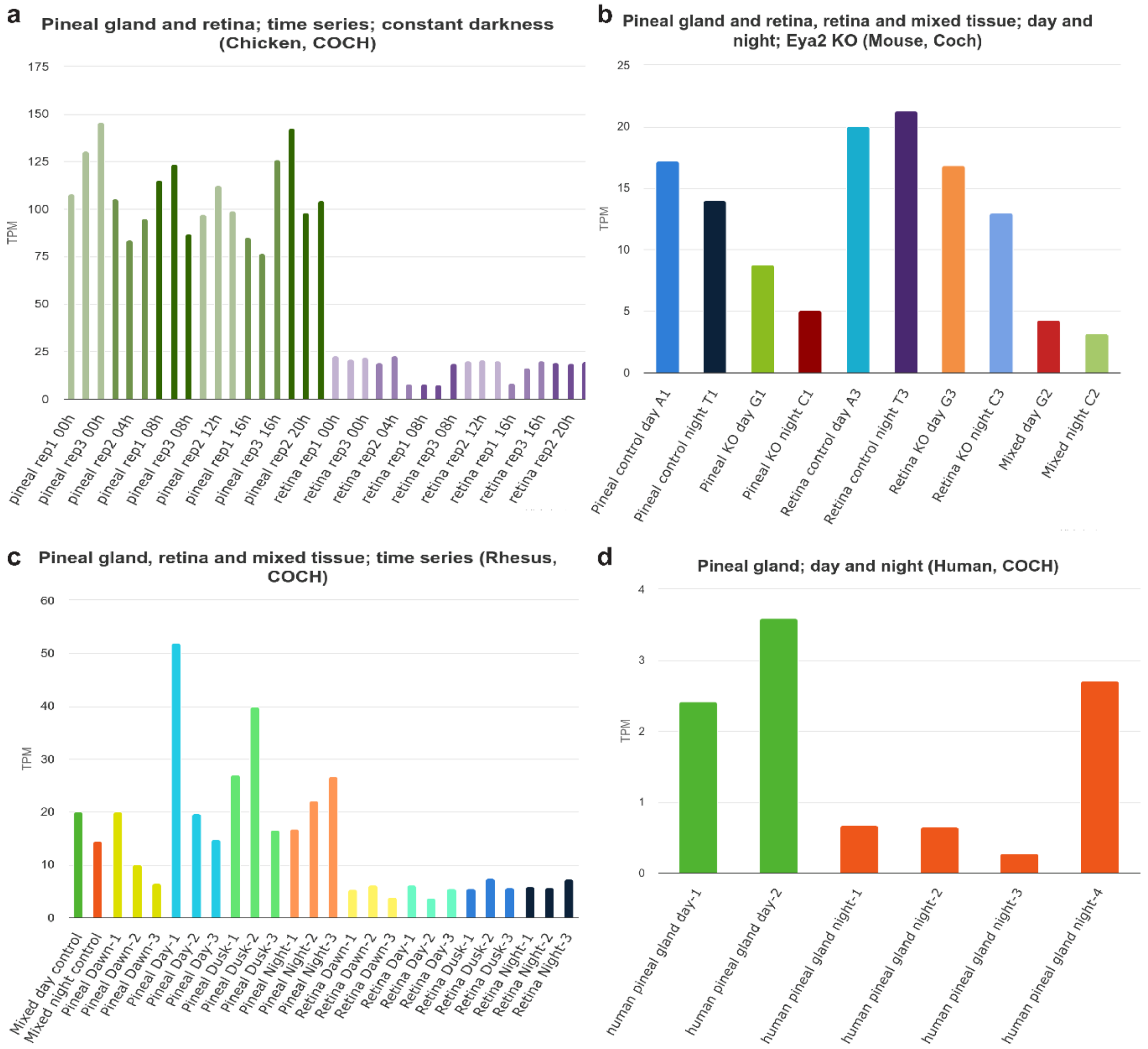


**Figure S8.** ***Cochlin*** **expression in the pineal glands of other vertebrates.** Data from the chicken (a), mouse (b) rhesus monkey (c) and humans (d) (4).

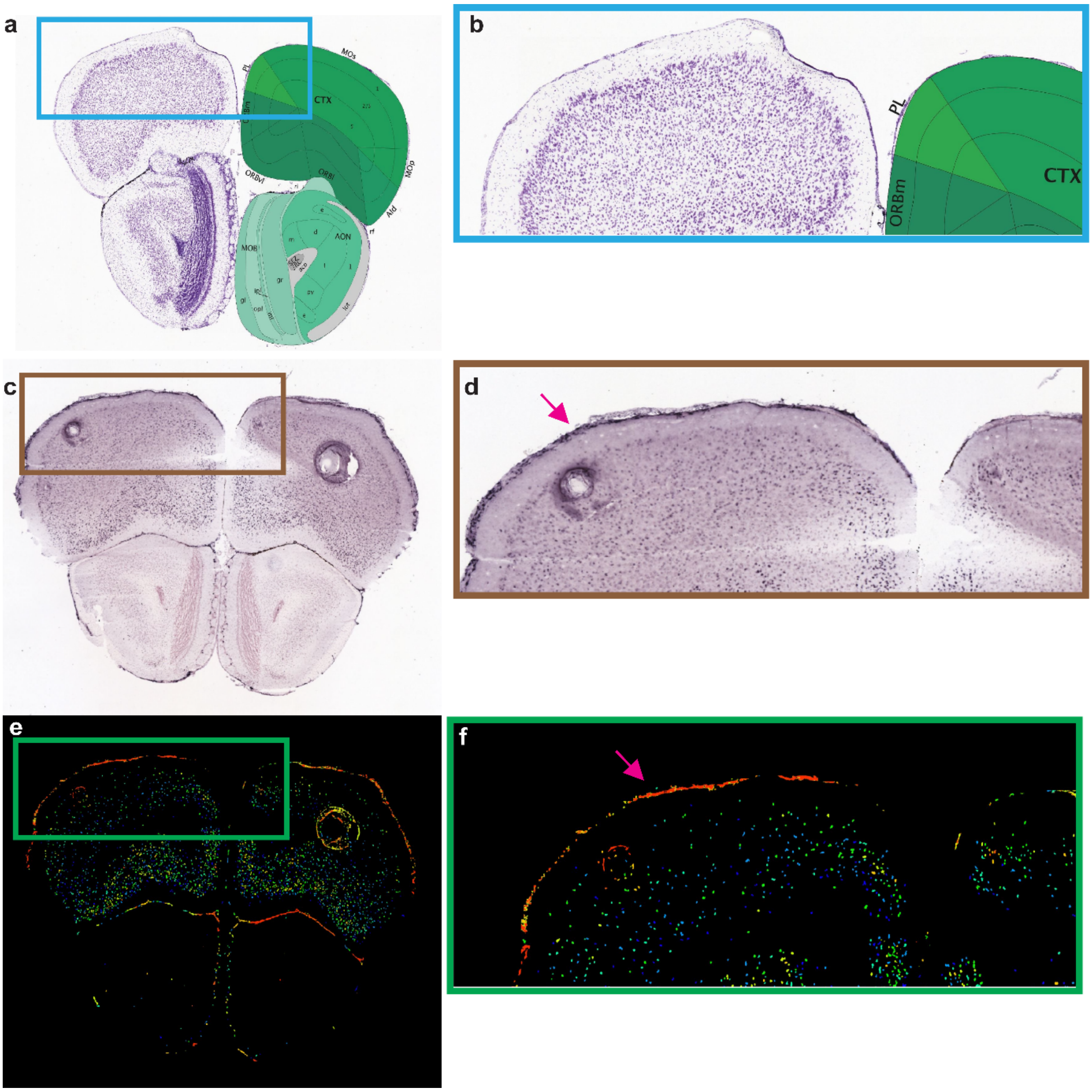


**Figure S9: *Cochlin* expression in the adult mouse meninges. a.** Nissl (left) and anatomical annotations (right) from the Allen Mouse Brain Atlas and Allen Reference Atlas – Mouse Brain, at the same region as (c) and (e). **b.** Anatomical annotations of zoomed out inset from panel a. Ctx - cortex, ORBm - Orbital area medial part, PL- Prelimbic area. Allen Reference Atlas – Mouse Brain (5). **c.** Expression of *Coch* in adult mouse brain (ISH). Allen Mouse Brain Atlas, mouse.brain-map.org/experiment/show/71717614 in the forebrain (5–8). **d.** Zoomed in inset from panel c. Magenta arrow: leptomeningeal staining. **e.** Expression heatmap of the section presented in panel c. **f.** Zoomed in inset from panel e. Magenta arrow: high expression in the leptomeninges.

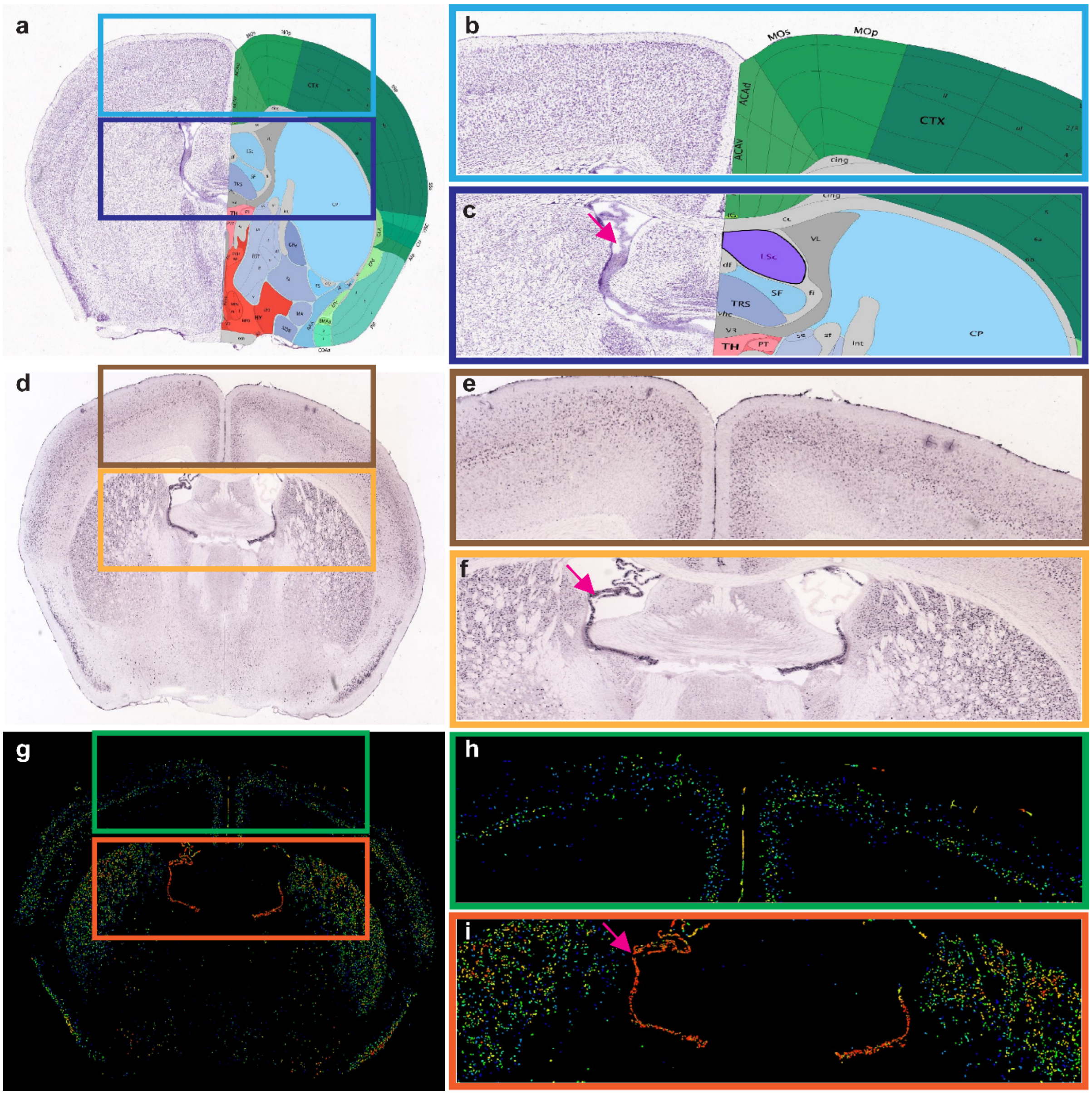


**Figure S10: *Cochlin* expression in the adult mouse brain. a.** Nissl (left) and anatomical annotations (right) from the Allen Mouse Brain Atlas and Allen Reference Atlas – Mouse Brain, at the same slice position as (d) and (g). **b, c.** Anatomical annotations of zoomed out inset from (a)- Ctx- cortex, ACAv- Anterior cingulate area -ventral, ACAd- Anterior cingulate area – dorsal, MOp- Primary motor area, Mos- Secondary motor area, df – dorsal fornix, VL – lateral ventricle, CP-Caudoputamen. Allen Reference Atlas – Mouse Brain (5). **d.** Expression of *Coch* in adult mouse brain (ISH). Allen Mouse Brain Atlas, mouse.brain-map.org/experiment/show/71717614 in the forebrain at the level of the septal nuclei (5–8). **e, f.** Zoomed in insets from panel d. **g.** Expression heatmap of the section presented in panel c. **h, i.** Zoomed in inset from panel e. Magenta arrows in panels f and i: high expression in the choroid plexus of VL.

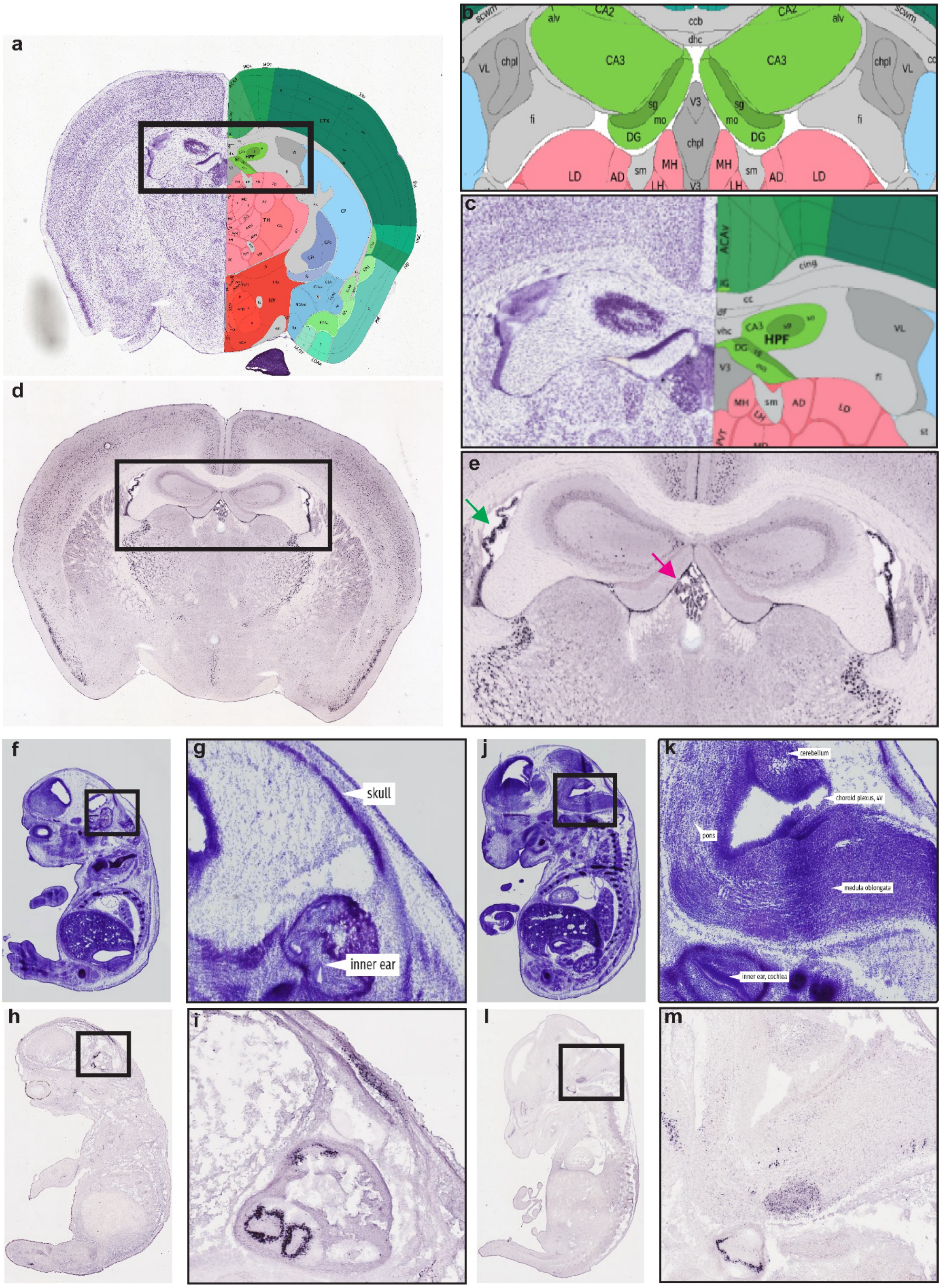

**Figure S11: *Cochlin* expression in the choroid plexus. a.** Nissl (left) and anatomical annotations (right) from the Allen Mouse Brain Atlas and Allen Reference Atlas – Mouse Brain, at the same slice position as panels b, c & d. **b.** Anatomical annotations of zoomed out inset from (a)- chpl-choroid plexus, VL-lateral ventricle, V3- third ventricle. Allen Reference Atlas – Mouse Brain (5). **c.** Nissl (left) and anatomical annotations(right) of zoomed in inset from panel a showing the structure of choroid plexus in the lateral and third ventricle. Allen Mouse Brain Atlas and Allen Reference Atlas – Mouse Brain (7). **d.** Expression of *Coch* in adult mouse brain (ISH). Allen Mouse Brain Atlas, mouse.brain-map.org/experiment/show/71717614 (5–8). **e.** Zoomed in inset from (d). green arrow- choroid plexus in the lateral ventricle, magenta arrow- choroid plexus of the third ventricle. **f-j.** Reference atlas for embryonic day 14.5 (E14.5) with Thioninacetate (Nissl-method) stain available on Genepaint (9). **g.** Zoomed in inset from (f) showing location of the inner ear and skull. **k.** Zoomed in inset from (j) showing location of the fourth ventricle (4V) choroid plexus. **h, l.** ISH for stained *Coch* in E14.5 mouse as available from https://gp3.mpg.de/results/Coch (9). **i.** Zoomed in inset from (h) showing staining of the inner ear, which serves as a positive control. **m.** Zoomed in inset from (l) showing the anatomical regions near the choroid plexus of the fourth ventricle.

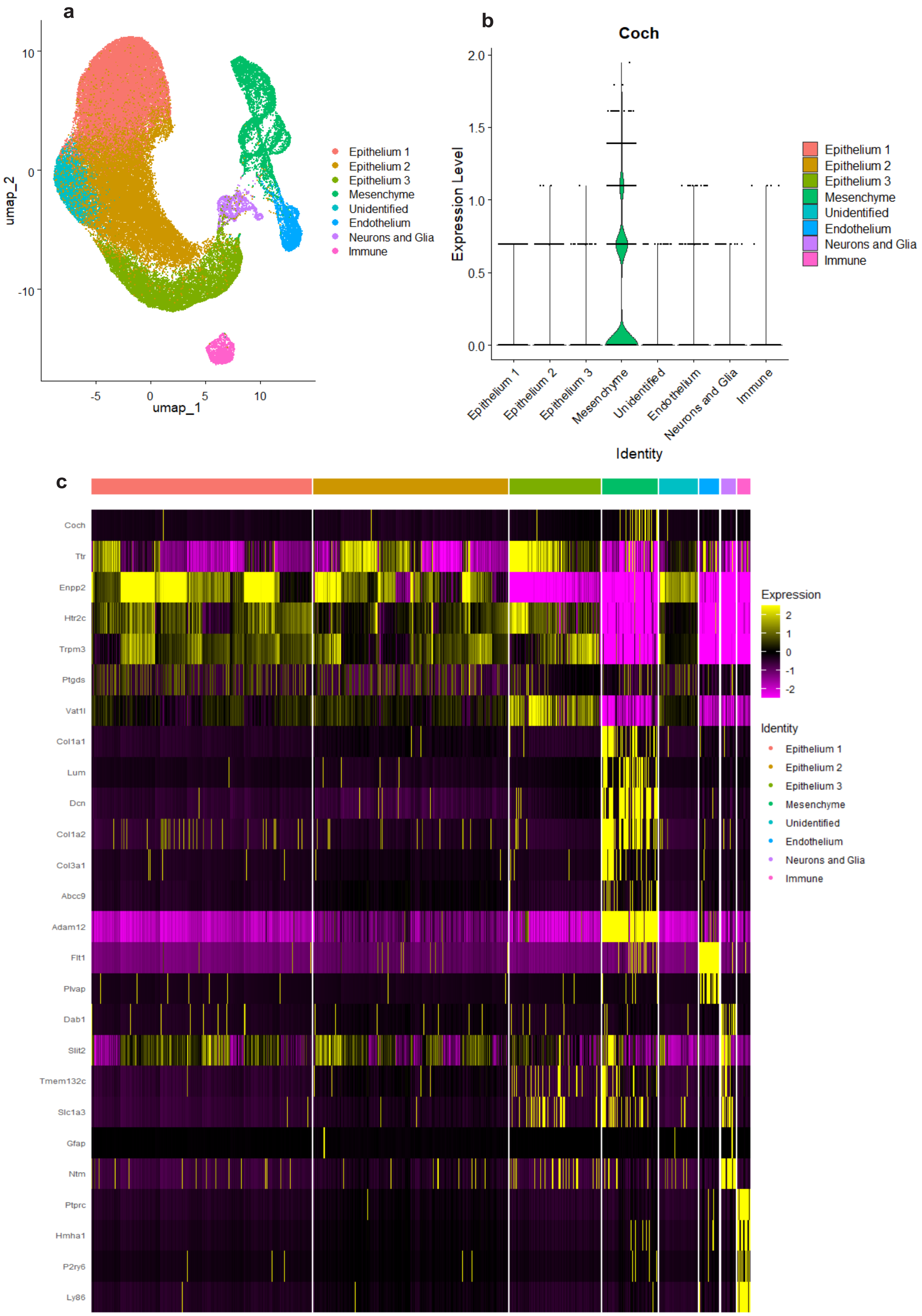


**Figure S12. *Coch* expression in murine choroid plexus. a.** UMAP of the reanalysed murine choroid plexus snRNA dataset (10). **b.** Violin plot of *Coch* expression as the clustering described in (a). **c.** Heatmap with selected markers used for cluster annotation along with *Coch*.

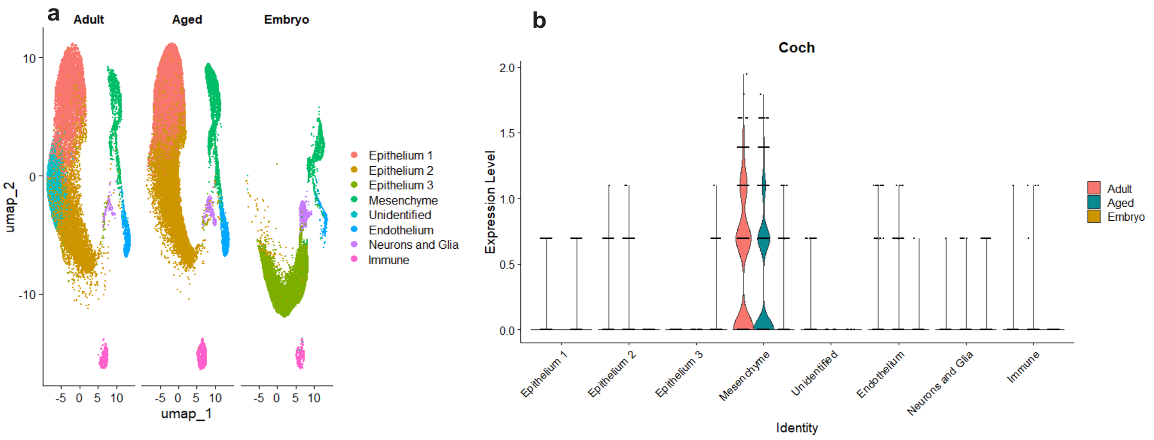


**Figure S13. *Coch* expression the mouse choroid plexus at different ages. a.** UMAP of the reanalysed murine choroid plexus snRNA dataset split by the age of the samples (10). **b.** Violin plot of *Coch* expression, split by age of the samples and clustered as described in Figure S9a.

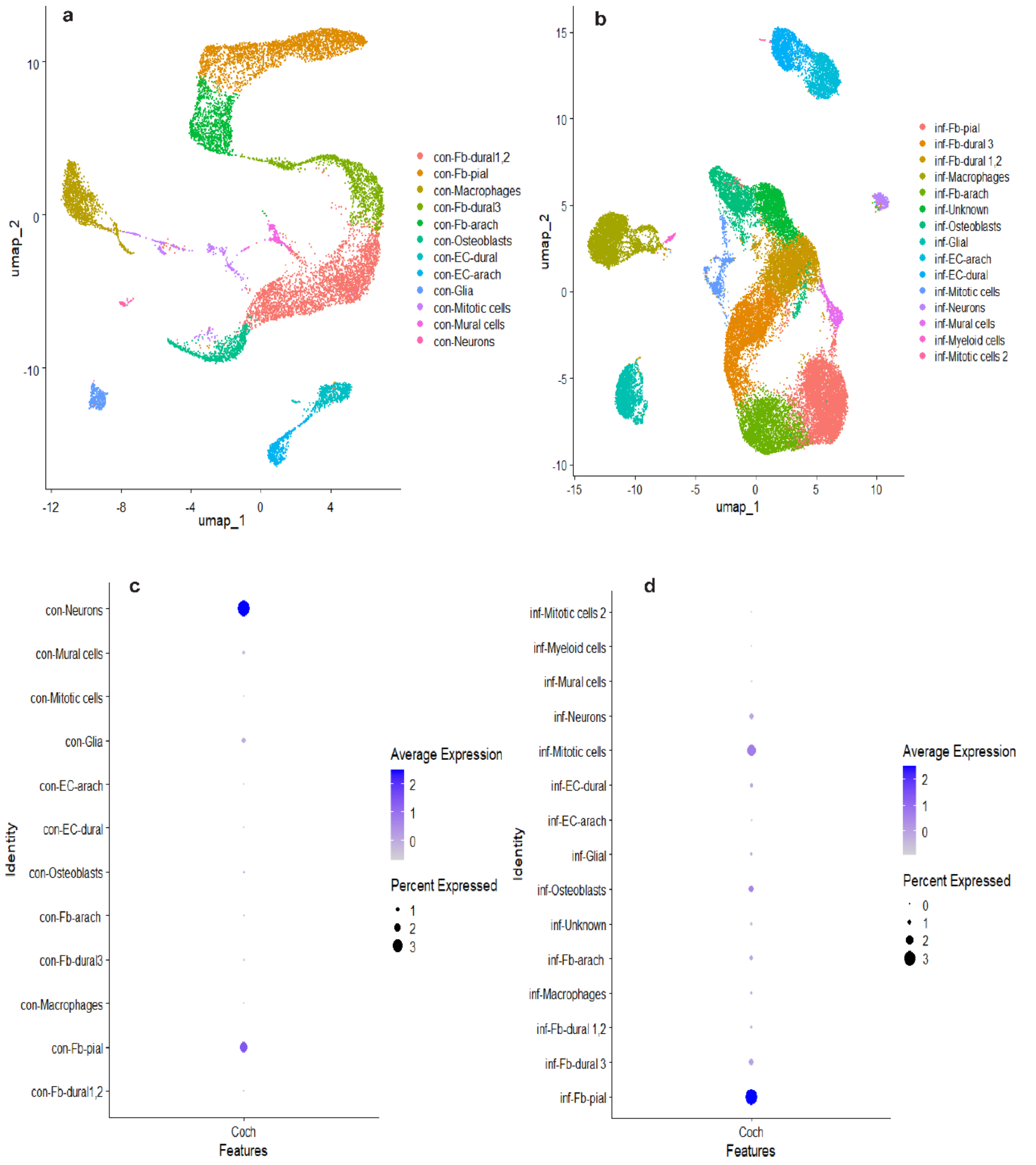


**Figure S14. Murine meningeal fibroblast expression of *Coch* in early post-natal mice with and without *E.coli* meningitis . a-b.** UMAPs of the integrated Seurat objects of meningeal cells of control (a) and infected (b) mice, based on reanalysis of GSE221678 (11). **c-d.** Dotplot for *Coch* expression in the integrated control (c) and infected datasets (d). con – control, inf- infected, Fb- fibroblasts, arach- arachnoid, EC- endothelial cells.

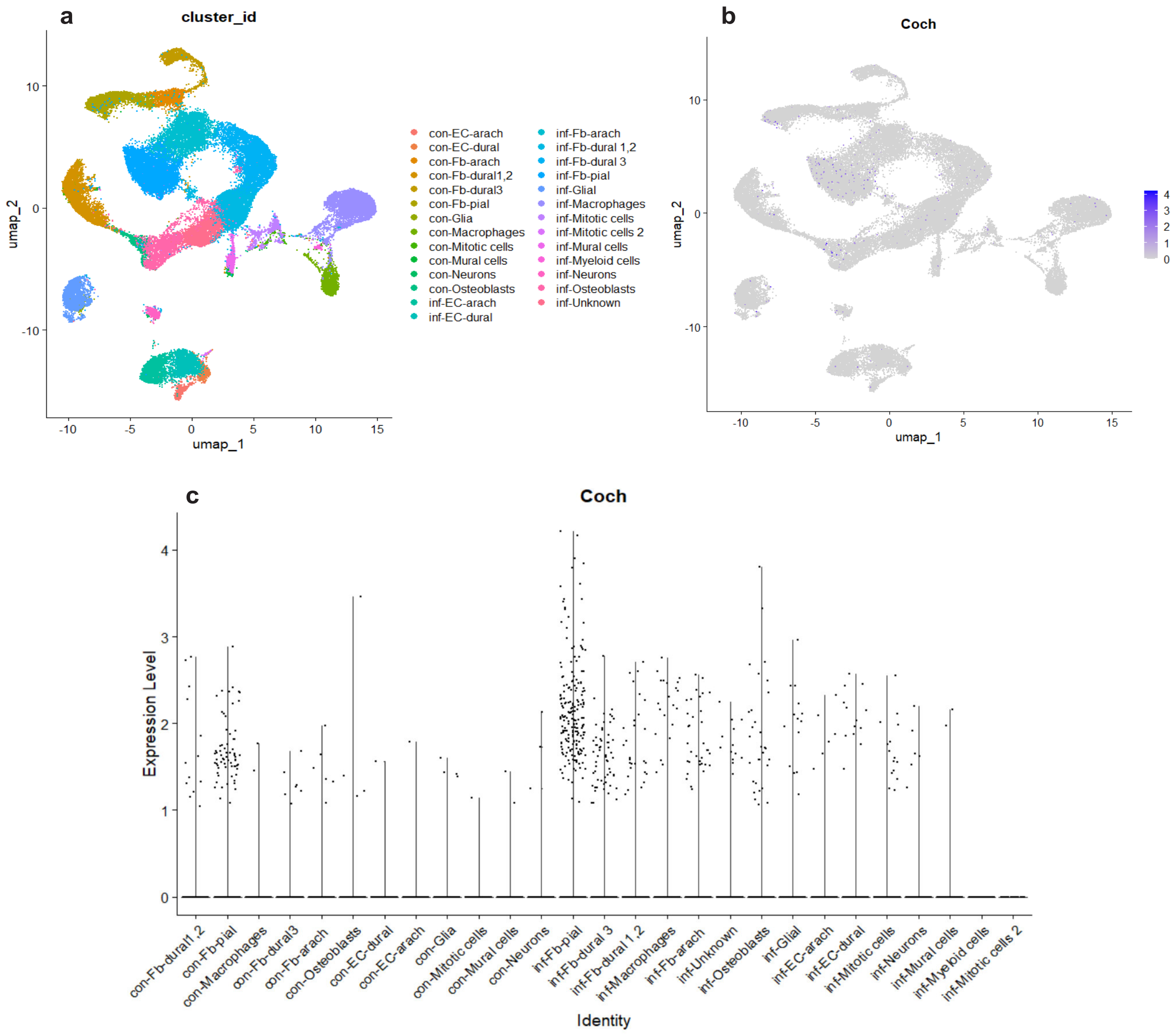


**Figure S15. Murine meningeal expression of *Coch* increases in early post-natal mice with *E.coli* meningitis. a.** UMAP of the merged Seurat object of meningeal cells of control and infected mice based on reanalysis of GSE221678 (11). **b.** Feature plot for *Coch* expression in the merged Seurat object. **c.** Violin plot for *Coch* expression in the merged Seurat object. con – control, inf- infected, Fb- fibroblasts, arach- arachnoid, EC- endothelial cells.

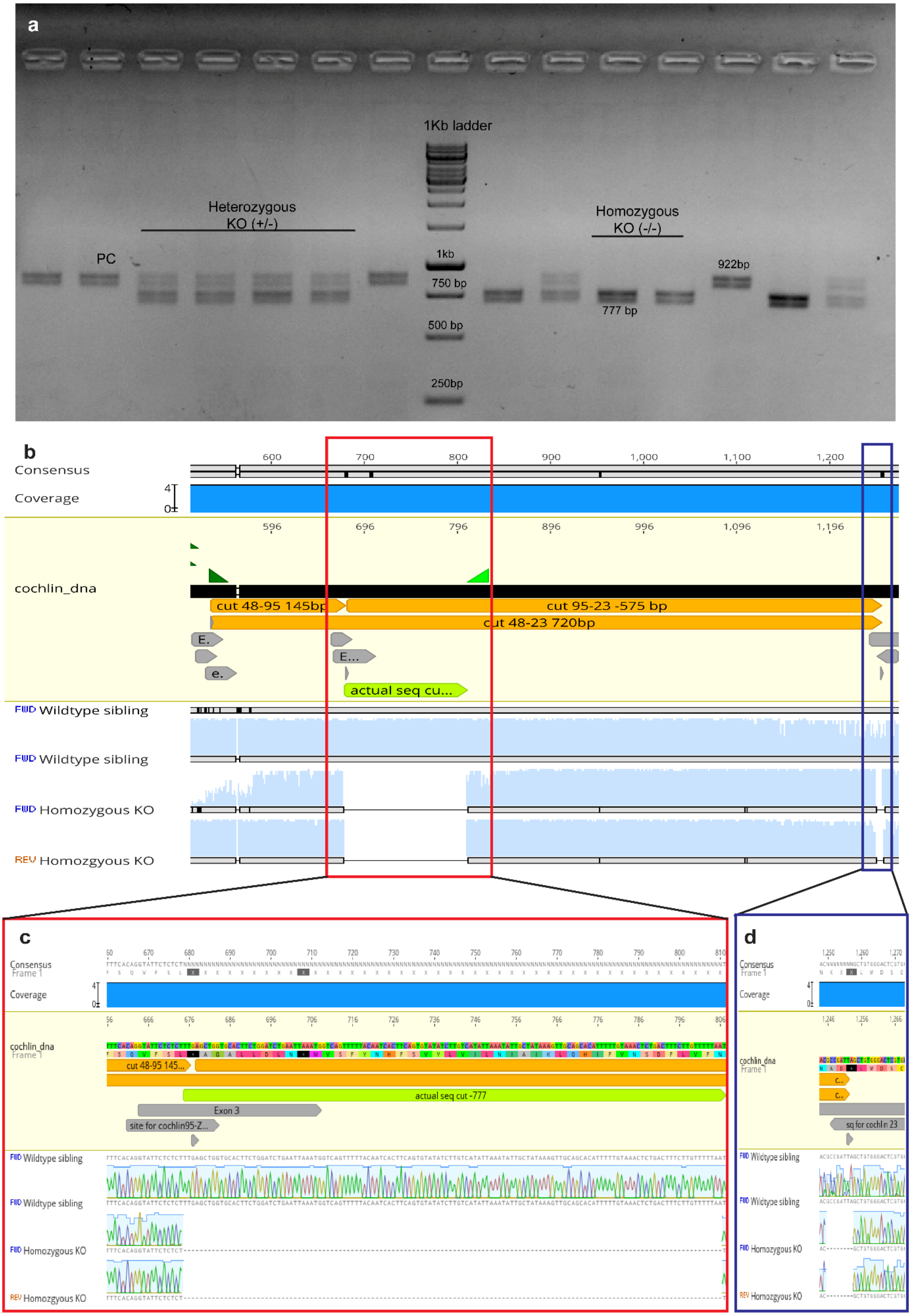
a
1Kb ladder
Heterozygous
KO (+/-)
PC
Homozygous
KO (-/-)
922bp
1kb
750 bp
500 bp
250bp
777 bp
b
Consensus
Coverage
cochlin_dna
cut 48-95 145bp
cut 95-23 -575 bp
cut 48-23 720bp
actual seq cu...
FWD Wildtype sibling
FWD Wildtype sibling
FWD Homozygous KO
REV Homozygous KO
c
Consensus
Frame 1
Coverage
cochlin_dna
cut 48-95 145...
actual seq cut -777
Exon 3
site for cochlin95-Z...
FWD Wildtype sibling
FWD Wildtype sibling
FWD Homozygous KO
REV Homozygous KO
d
Consensus
Frame 1
Coverage
cochlin_dna
Frame 1
sq for cochlin 23
FWD Wildtype sibling
FWD Wildtype sibling
FWD Homozygous KO
REV Homozygous KO

**Figure S16. Sequencing of the *cochlin* gene in *coch*$^{lkcsj1}$. a.** Gel electrophoresis image of genotyping PCR of offspring derived from in-cross of F1 *cochlin* heterozygous mutants. PC-Positive control. **b.** Overview of the sequencing results of two homozygous mutant (-/-) and one wt (+/+) sibling. **c.** Deletion at the cut site for cochlin95 gRNA. **d.** Deletion at the cut site for cochlin23 gRNA.

**TableS1.** Sex of adult HCR-ISH samples

| Genes | Female | Male |
|---|---|---|
| *rgra* | *n* = 4 (Represented in the FigS11 q-t) | *n* = 2 |

7. ISH Data :: Allen Brain Atlas: Mouse Brain [Internet]. [cited 2024 Jul 31]. Available from:

https://mouse.brain-map.org/

8. Lein ES, Hawrylycz MJ, Ao N, Ayres M, Bensinger A, Bernard A, et al. Genome-wide atlas of gene expression in the adult mouse brain. Nature. 2007 Jan;445(7124):168–76. doi:10.1038/nature05453

9. Genepaint - Home of High Resolution Gene Expression Data [Internet]. [cited 2024 Jul 31]. Available from: https://gp3.mpg.de/

10. Dani N, Herbst RH, McCabe C, Green GS, Kaiser K, Head JP, et al. A cellular and spatial map of the choroid plexus across brain ventricles and ages. Cell. 2021 May;184(11):3056-3074.e21. doi:10.1016/j.cell.2021.04.003

11. Wang J, Rattner A, Nathans J. Bacterial meningitis in the early postnatal mouse studied at single-cell resolution. eLife. 2023 Jun 15;12:e86130. doi:10.7554/eLife.86130
